# Covalent Inhibition of the Human Papillomavirus Type 16 E6 Protein Restores p53 and Suppresses HPV-Driven Tumorigenesis

**DOI:** 10.1101/2025.08.24.671874

**Authors:** Anne Rietz, Lokesh Kumari, Ankeeta Koirala, Stephane Pelletier, Zhijian Lu, Elliot J. Androphy

**Affiliations:** Department of Dermatology, Indiana University School of Medicine, Indianapolis, IN, USA; Department of Medical and Molecular Genetics, Indiana University School of Medicine. Indianapolis, IN, USA; Kovina Therapeutics Inc. Indianapolis, IN, USA

## Abstract

High-risk human papillomavirus (HPV) infections are the etiology of approximately 5% of all cancers worldwide, including cervical, anal, and oropharyngeal malignancies. HPV E6 is a multifunctional oncoprotein that drives tumorigenesis and is best known for bridging the ubiquitin ligase E6AP (UBE3A) and p53 into a complex that leads to proteasome mediated destruction of p53. We developed small molecule inhibitors that covalently bind to cysteine-51 (Cys-51) in HPV16 E6. In HPV16-positive cancer cells, these compounds increase p53 protein levels and activate p53-dependent transcriptional programs associated with apoptosis and senescence, resulting in reduced tumor cell viability. *In vivo*, E6 inhibition suppresses the growth of human HPV16 expressing cervical and oropharyngeal tumor cell lines in mice. The strategy of targeting a viral oncoprotein with a covalent inhibitor demonstrates a genotype-specific therapeutic strategy for HPV-associated cancers and premalignant infections, addressing a significant unmet need in current treatment options.

## INTRODUCTION

There is no specifically antiviral treatment for the many millions of people afflicted with human papillomaviruses (HPV) infections. While infections of the uterine cervix, anus, genitalia and oropharynx are very common, most remain benign and spontaneously resolve. However, persistent infections with specific ‘high risk’ (HR) HPV genotypes can progress to invasive and metastatic cancers. It is estimated HPVs are the etiologic agent of ∼5% of all cancers in the world (1, 2). These epithelial malignancies evolve over several years to decades from benign precursor pathologic stages at the site of infection. This timeline offers a realistic interval for medical intervention. Development of an effective specifically antiviral therapy is hampered by the fact that HPVs have limited early protein targets and only its E1 protein has the traditionally targeted enzymatic activity, which has not been successfully discriminated from host ATPases.

The E6 protein is necessary for stable viral genome replication and episomal maintenance, and for HR HPV transformation of keratinocytes (3–5). E6 has multiple activities and has been reported to bind more than a dozen cellular proteins. The two best characterized are the HECT domain ubiquitin ligase E6AP (UBE3A) and the tumor suppressor p53. E6 does not bind to p53 in the absence of E6AP and E6AP does not bind to p53 without E6. In this complex, E6AP transfers ubiquitin onto p53 and mediates its destruction by the proteasome (6) (Figure 1). Both HPV16 E6 and E7 are consistently expressed in HPV-induced tumors, which is relevant to our approach as HR HPV E7 alone activates p53 (7, 8). RNAi-mediated knockdown or knockout of E6 restores wild-type p53 protein levels and activity, leading to growth arrest or apoptosis of HPV cervical cancer cells (9–15). A recent study of siRNA mediated depletion of E6AP demonstrated dramatic induction of cell senescence that depended on E7 mediated inhibition of the pRb tumor suppressor and the p53/p21 pathways (16). Our chemical approach to inactivate E6 does not interfere with expression of the E7 protein.

**Figure 1.**
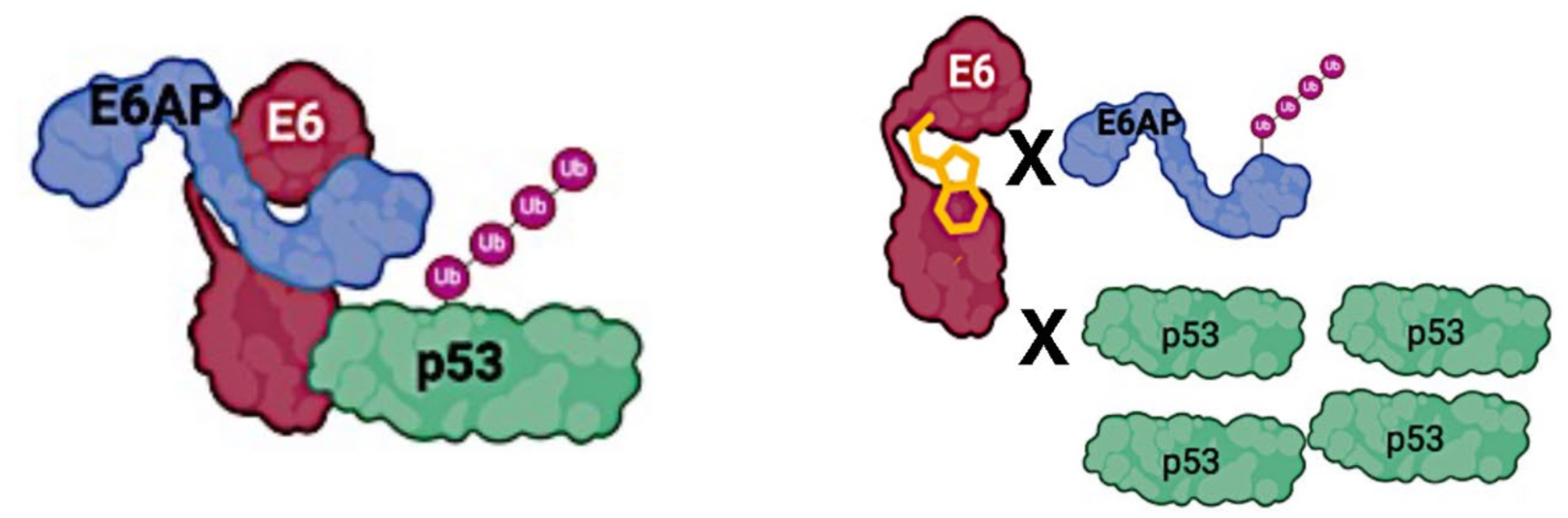
Conceptualization of compound blocking the interactions of E6 with E6AP and p53. E6 binding to the ubiquitin (Ub) ligase E6AP stabilizes a p53 binding surface on E6, leading to the recruitment of p53 by the E6•E6AP complex. E6AP transfers Ub to the p53 DNA, leading to p53 degradation by the proteasome. Compound depicted in yellow covalently binds to a specific cysteine in E6 and block E6AP and p53 association, thereby preventing p53 ubiquitination and degradation.

An α-helical peptide in E6AP containing an LxxLL motif, where L is leucine, docks into a well-defined pocket of E6 (17–22). Travé and co-workers solved the crystal structure of a complex that includes maltose binding protein (MBP) fused to an E6AP peptide and HPV16 E6 (PDB code 4GIZ) (19). This group also solved the trimeric crystal structure of HPV16 E6 in complex with the MBP-E6AP peptide and the core domain of p53 (PDB code 4XR8) (21). Association with E6AP exposes a large p53 interaction surface on E6 and generates the E6•E6AP•p53 trimeric complex (21, 23). NMR studies of free E6 in solution indicated that its two zinc binding domains can adopt various positions relative to each other and that the connecting helix that separates these is not fully stabilized (19). Its association with the LxxLL peptide of E6AP stabilizes E6 into a single conformation in which the connecting helix becomes rigid and the two zinc-binding domains form a single coherent unit around the bound peptide. In this conformation, both zinc binding domains contribute to form a large p53 interaction surface. This E6AP binding pocket also serves as a ‘hot spot’ for association with a diverse set of cellular proteins that encode an LxxLL motif including E6BP, paxillin, E6TP1, CBP/p300, tumor necrosis factor receptor 1, Mcm7, Tyk2, IRF-3, pro-caspase-8 and hADA3 [reviewed in (19, 24, 25)]. Here we describe the biological effects of small molecules designed to irreversibly bind to and inactivate the HPV16 E6 protein and thereby restore p53 levels and activities with the expectation that other functions of E6 would also be blocked (Figure 1).

## RESULTS

### Activities of Compounds that Specifically Inhibit HPV16 E6 Mediate p53 Degradation

Screening and biochemical characterization of a series of compounds that specifically bind to and form a covalent bond with cysteine at position 51 (Cys-51) proximal to the E6AP binding pocket of the HPV16 E6 protein will be reported elsewhere. Some publications refer to this as cysteine 58 due to an upstream ATG codon that adds 7 amino acids onto the E6 reading frame. Here we demonstrate the biological effects of two compounds: KTI-218 and KTI-240 (Figures 2A, B). To test for activity in live cells, we constructed a luciferase based ‘gain of signal’ reporter assay to measure inhibition of E6•E6AP-mediated p53 degradation. Wild-type p53 was cloned in frame with renilla luciferase (R-Luc). The E6•E6AP complex binds the p53 moiety of the fusion protein and induces degradation of the chimeric protein (26, 27). These DNA constructs were transfected into HPV16 expressing human cervical cancer derived cell line SiHa (ATCC-HTB35™), which was selected because levels of E6 and E7 are directed by the native HPV16 promoter. RPE-1 cells (hTERT immortalized epithelial cells that express wild-type p53; HPV-negative) expressing the identical p53-R-Luc reporters were also isolated. The E6 mediated pathway for degradation of p53 in the cervical cell lines was validated using siRNAs to inhibit expression of E6E7, E6 and E6AP, which resulted in increased p53-R-Luc signal, whereas scrambled control siRNA did not (Supplementary Figure 1A). Following incubation of SiHa cells in media with KTI-218 or KTI-240, p53-luc increased by 8- and 14-fold with an EC50 of 4.2 ± 0.1 µM and 2.3 ± 0.1 µM, respectively, without changing p53-luc activity in the control RPE-1 reporter cells (Figures2A, B). To investigate the contribution of the covalent group, we prepared KTI-239, a non-covalent analog of KTI-218. KTI-239 induced p53-luc activity at a much lesser extent than KTI-218 in SiHa-luc cells without affecting Luc activity in RPE-1 cells (Figures 2C, D).

**Figure 2.**
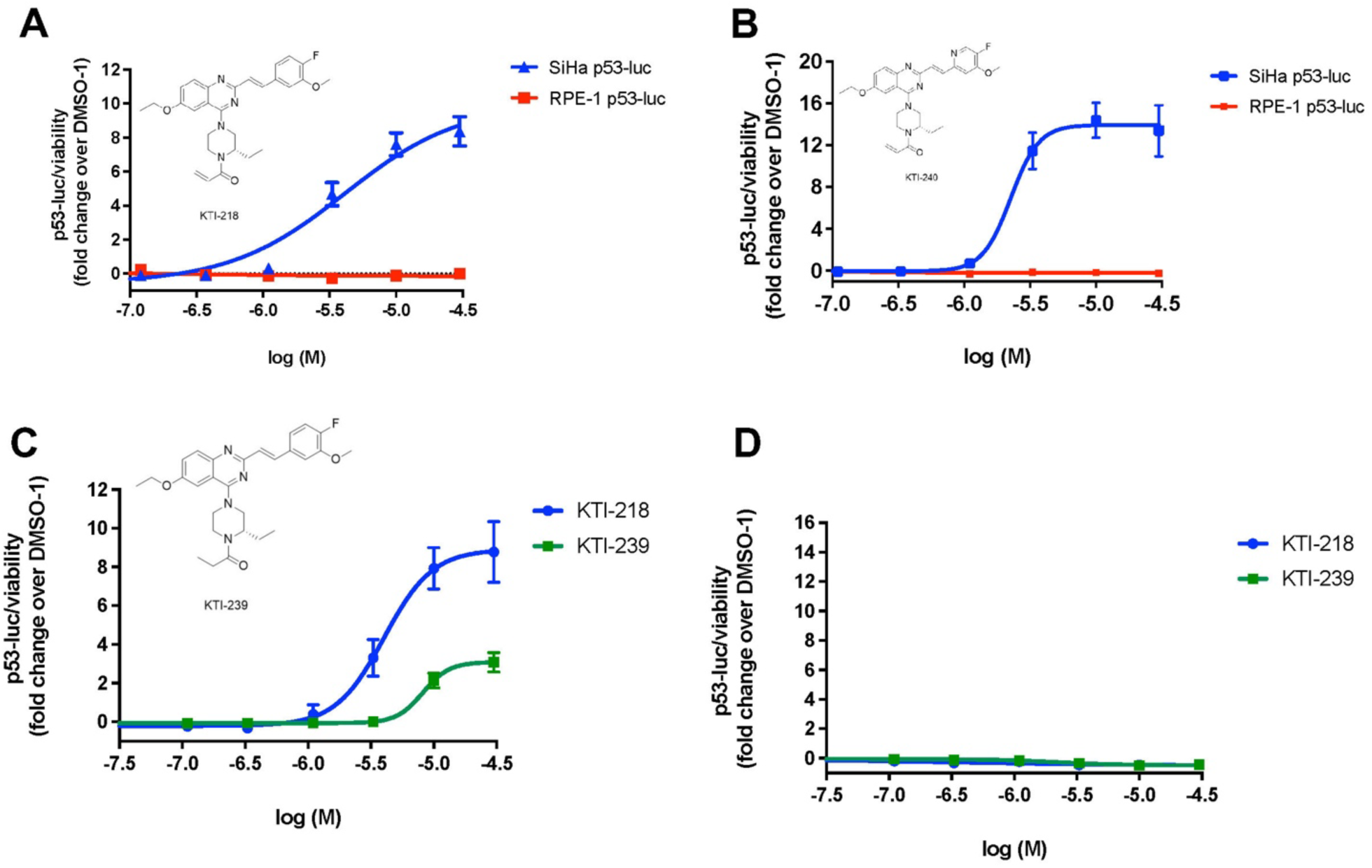
E6 inhibition in cell-based reporter assay. A stable clone of HPV-16 E6/ E7 expressing human cervical cancer SiHa cells with a p53-Luciferase (p53-Luc) reporter and a clone of HPV-negative RPE-1 cells expressing the same p53-Luc reporter were incubated with DMSO (0.02% (v/v)) or increasing concentrations of compounds added to the media. SiHa-P53-Luc cells (red) and RPE-luc cells (blue) were treated with increasing concentrations of **(A)** KTI-218 or **(B)** KT-240 for 24 hrs. To study the contribution of the covalent warhead, we prepared, KTI-239, an analog of KTI-218 that lacks the acrylamide warhead and tested in SiHa-P53-Luc **(C)** and RPE-luc cells **(D**), respectively. Structures of each compound is shown in the insets. Luciferase and cell viability were measured after 24 hours. Luc signal was normalized to cell viability and p53-Luc induction is expressed as fold-change over DMSO control. Data is expressed as S.E.M and each experiment was completed at least three independent times (n ≥ 3).

Confirmation of E6 inhibition was performed by evaluating cellular p53 protein levels following exposure to compound in culture media. Western blots showed robust p53 elevation in HPV16-positive SiHa cells (Figures 3A, B) but not in p53 wild type RPE-1 cells (Figure 3C, D), implying this induction is not due to off-target genotoxic stress. Positive control etoposide induced p53 in both cell lines, confirming an intact p53 pathway. KTI-239 lacks the acrylamide warhead in KTI-218 and did not produce a rise in p53 levels. Increased levels of p53 protein in SiHa cells were detectable by 6 hours in culture with KTI-218 (Figures 3E, F). This rapid response aligns with E6’s ∼3-hour half-life (28), highlighting the compound’s potency against nascent viral protein.

**Figure 3.**
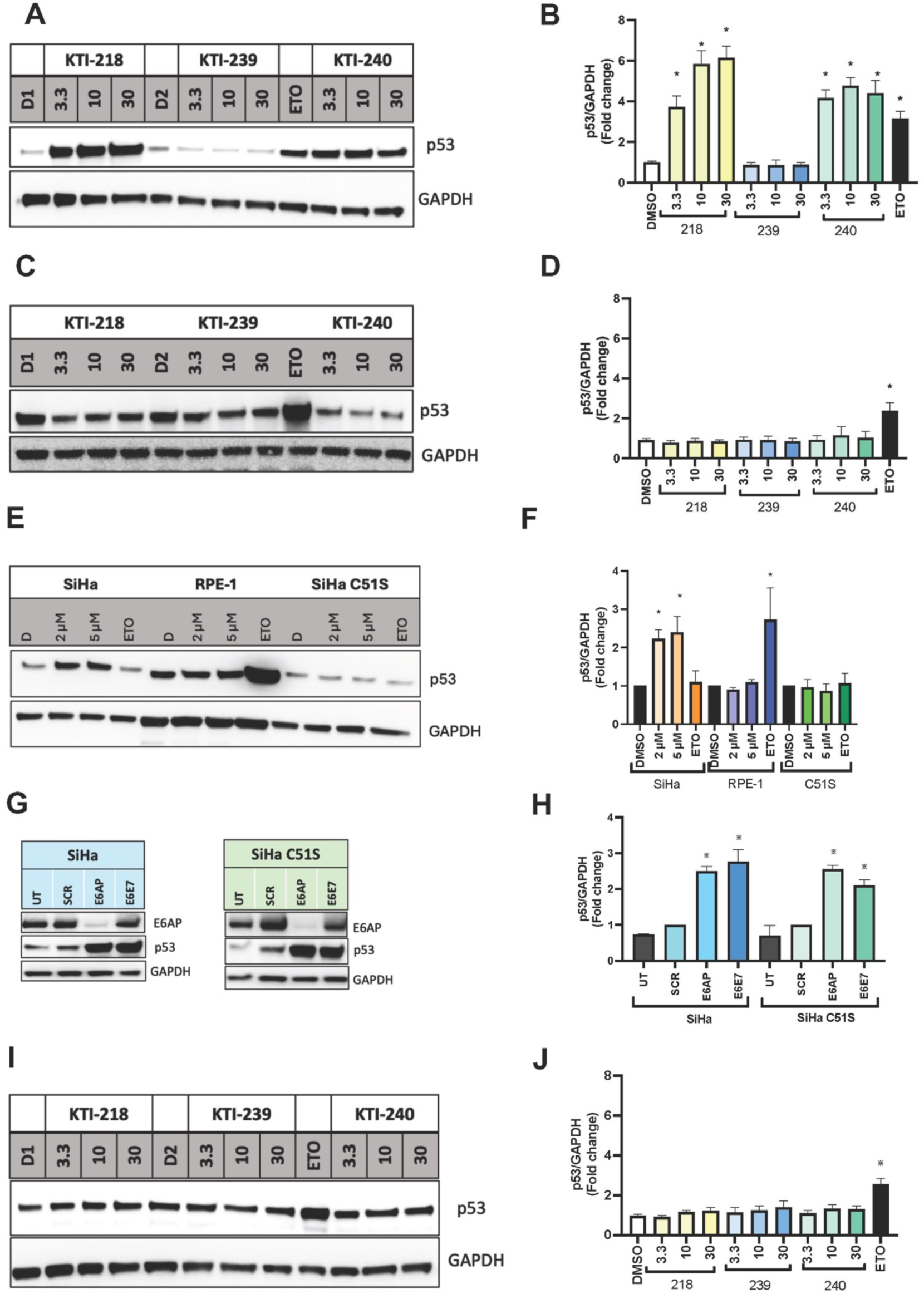
Compounds selectively increase p53 protein and require cysteine 51 in HPV16 E6. **(A, B)** P53 protein levels in SiHa and **(C, D**) RPE-1 cells cultured in KTI-218, KTI-239 and KTI-240 for 24 hours. **(E, F)** KTI-218 robustly increased p53 protein levels with 2-fold within 6 hours of incubation with in SiHa cells but not in RPE-1 or SiHa C51S cells. **G, H**) siRNA knockdown of E6AP and 16E6E7 result in comparable p53 inductions in both SiHa and SiHa C51S cells, in which the codon for cysteine 51 was converted to serine. SCR–scrambled RNAi control. UT – untreated cells. **(I, J)** SiHa C51S cells were treated with KTI-218, −239, and −240 for 24 hours and did not induce p53 protein levels. ETO – etoposide. D1, D2 – independent DMSO controls. Protein expression was normalized to GAPDH and expressed as fold-change. Data is expressed as S.E.M and analyzed using were analyzed using one-way ANOVA with Dunnett’s or Bonferroni post hoc analysis. Each experiment was completed at least three independent times (n ≥ 3; * p< 0.05).

To provide evidence for the requirement of covalent bonding to Cys 51 for E6 inhibition, CRISPR mediated mutagenesis was used to convert the codon for cysteine 51 in HPV16 E6 to serine. This was possible because unlike most human-derived HPV cancer cells that have tens to hundreds of copies HPV16, SiHa cells have only one or two integrated copies. We isolated a C51S clone and confirmed it was derived from the parental SiHa cell line by autosomal STR profiling and that the DNA sequences of the E6 region were otherwise preserved (Supplementary Figures 2A, B). The C51S SiHa cells proliferated in culture similar to the parental cells. Levels of p53 were comparable to the parental SiHa line (Supplementary Figure 2C), which was expected, for while high-risk HPV E6 proteins bind to E6AP, the targeted cysteine is exclusive to HR HPV types 16, 35, 45, and 68 and is not present in other oncogenic HPV types. RNAi mediated depletion of E6AP or HPV16 E6 resulted in increased p53 levels similar to wild type SiHa cells (Figure 3G, H), demonstrating the E6•E6AP pathway for p53 degradation is intact in the CRISPR-edited SiHa cells and that cysteine 51 can be functionally replaced by serine (29). Importantly, incubation of C51S SiHa with KTI-218 or KTI-240 did not increase p53 levels, while positive control etoposide did (Figures 3 I, J). These results infer that the observed p53 inductions depend on covalent binding of the compound to cysteine 51 in HPV16 E6 and are not due to a consequence of a covalent interaction with a cellular protein.

Levels of p53 protein also increased in HPV16 E6 expressing human oropharyngeal cancer (OPC) cancer derived cell lines such as UM-SCC-47 and UM-SCC-104 (Figures 4A, B) following compound exposure. Elevated levels of the p53 transcription target p21 protein were also detected (Figure 4A, B; Supplementary Figures 3A-D). We next questioned whether these compounds could inhibit E6 activity in the context of human keratinocytes that maintain episomal HPV16 genomes and express all early viral proteins. W12-E cells, which were derived from a woman with a pre-malignant cervical lesion, showed robust induction of p53 levels following incubation in media with KTI-218 and KTI-240 (Figure 4C). Etoposide was included as a control to induce DNA damage and p53, however in the majority of our experiments with HPV16 E6 expressing cells and as shown for the OPC line SCC-104, this did not induce p53, presumably due to overriding effect of E6 mediated p53 degradation.

**Figure 4.**
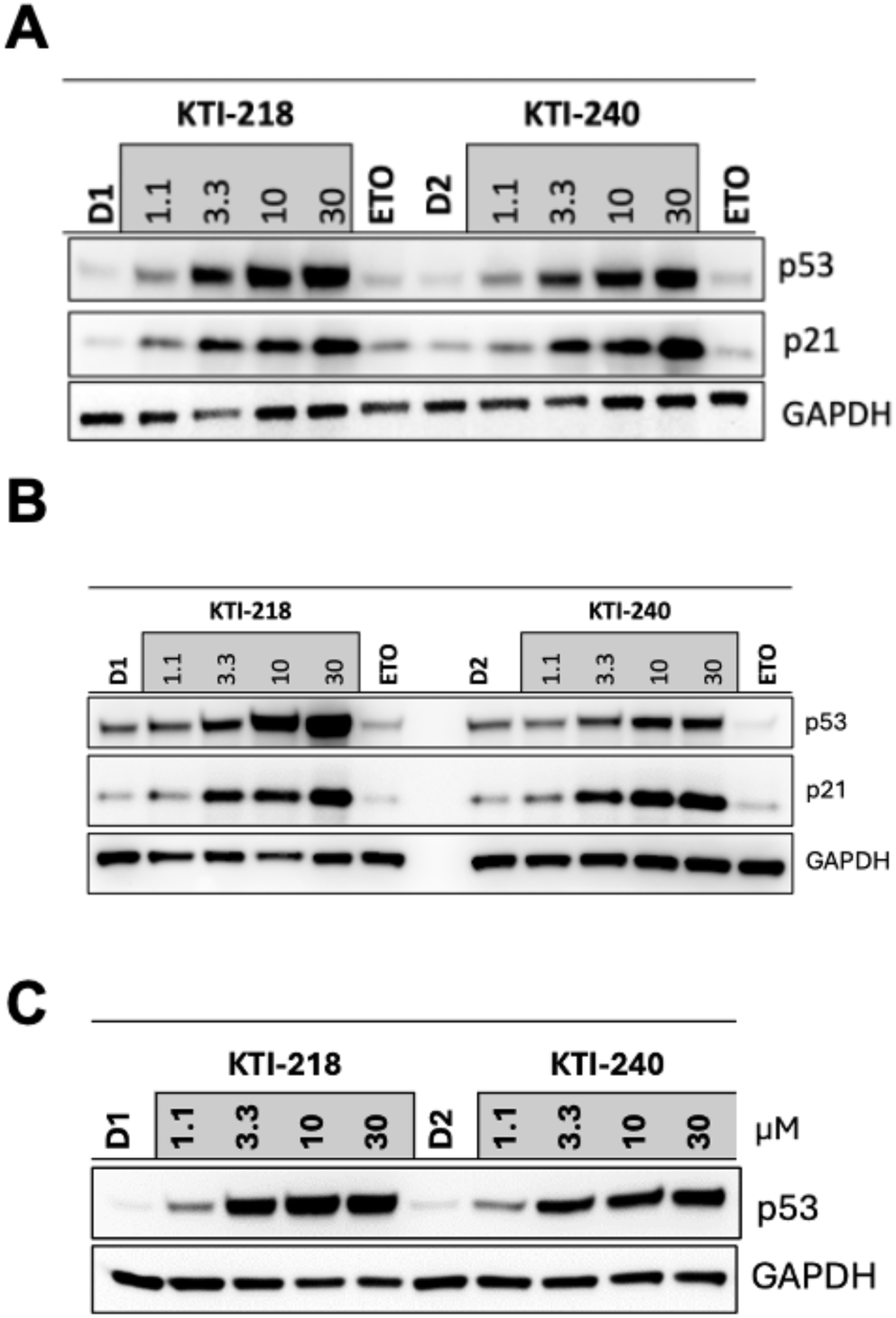
E6 inhibition increases p53 in HPV-16 E6 expressing human oropharyngeal cancer and cervical dysplasia cells. SCC-47 **(A)** and SCC-104 cells **(B)** were cultured with KTI-218, KT-240 or DMSO (D1, D2) for 24 hours, harvested and p53 and p21 protein levels were analyzed by immunoblot. Protein expression was normalized to GAPDH. In parallel, cells were treated with etoposide (ETO, 25 µM). **(C)** p53 levels in human cervical dysplasia cell line W12-E cultured in KTI-218, KT-240 or DMSO (D1, D2) for 24 hours.

### Transcriptomic profiling following E6 inhibition

We then sought to characterize the transcriptional program of HPV E6/E7 expressing tumor cells following E6 inhibition. The elevated levels of p53 protein are predicted to alter expression of its transcriptionally regulated target genes. While the C51S mutated SiHa cells did not show endogenous p53 induction following incubation in media containing KTI-218 and KTI-240, it is possible that some functions of E6, especially those pathways mediated by other binding partners of E6, could manifest and be differentiated from downstream effects of p53. The C-terminus of E6 binds to several PDZ proteins, and while this domain is distinct from the region of E6 targeted by our compounds, its interactions might also be affected. RNA from compound treated RPE-1 cells was included in these transcriptome studies to disclose potential off-target consequences in the absence of HPV E6. In this set of experiments, SiHa, SiHa C51S, and RPE-1 cells were cultured in 5 µM KTI-240 or DMSO for 16 hours. This time point was selected to avoid accumulation of non-viable cells, and we confirmed p21 protein was elevated within this time frame (Figure 9A). Total RNA from each of the 18 samples isolated from cultures harvested from three independent experiments was submitted to quality control analysis. Each sample had a RIN of >9 and mRNA was sequenced with an average depth of ∼50 x 10^6^ reads per sample. Parallel lysates were tested for p53 by Western blot and revealed a robust 2.2 ± 0.2 -fold induction of p53 (Figure 5A; Supplementary Figure 4A).

**Figure 5.**
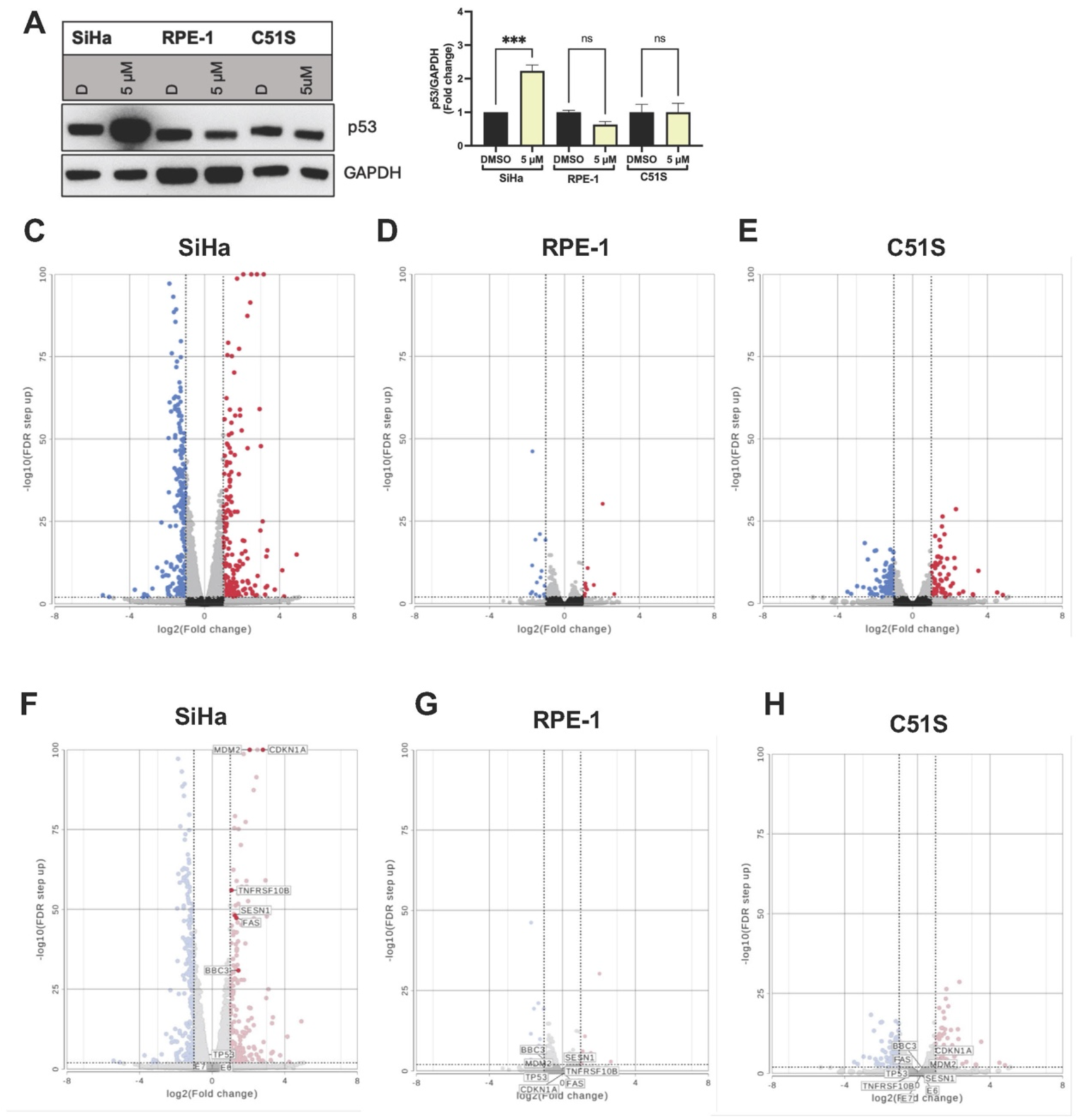
Transcriptional responses to E6 inhibition by KTI-240. SiHa, SiHa C51S, and RPE-1 cells were cultured in KTI-240 for 16 hours and RNA was extracted and sequenced from three independent experiments. **(A, B)**. Parallel cell cultures were analyzed for p53 protein levels by western blots showing a robust induction of p53. Data presented as S.E.M (n=3; * p<0.05). **(C-H)**. Differentially expressed genes were identified using DESeq2 analysis with a false discovery rate of <0.01 and a fold change of >2. Volcano plots showing significant transcriptional increases (right, red) and decreases (left, blue) comparing vehicle vs. KTI-240 treated cells. HPV-16 E6 and E7 mRNAs were detected and did not differ between WT and C51S SiHa cells lines. (**F, H**). TP53 mRNA was unchanged in all three cell lines upon KTI-240 treatment. Increased expression of p53 regulated genes occurred only in wild-type SiHa but not C51S or RPE-1 cells (**F, G, H**). p53 protein expression was normalized to GAPDH and expressed as fold-change. Data is expressed as S.E.M and was analyzed with one-way ANOVA with Dunnett’s or Bonferroni post hoc analysis. Each experiment was completed three independent times (n = 3; * p < 0.05).

The mRNA sequences of a total of ∼28,000 genes were identified in the native SiHa cells. Expression of E6, E7 and p53 mRNAs were unchanged following exposure to KTI-240 (Figure 5F; Supplementary Table 1). With a false discovery rate (FDR) of < 0.01, fold change (FC) of ≥ 2.0 and mean reads >5, expression of 197 genes increased, and 221 genes decreased following exposure of SiHa cells to KTI-240. Volcano plots are shown in Figure 5C with relevant genes indicated in Figure 5F. Supplementary Table 1 lists the differentially expressed genes (DEGs). Several p53 target genes such as p21(CDKN1A), MDM2, and Sestrin 1 (SESN1) increased after KTI-240 treatment (Figure 5F). p53 transcription targets that mediate apoptosis including FAS, Trail-r2 (TNFRSF10B), and PUMA (BBC3) are in this upregulated mRNA set. Some p53 responsive genes were minimally changed. For instance, the splicing factor Zmat3 increased but with an FC of 1.5 and FDR of <0.05 (30). The p53 target FoxM1 transcript decreased with KTI-240 treatment. FoxM1 is a pro-proliferative transcription factor that is repressed by p53 during the DNA damage response and is often over-expressed in cancers (31).

The basal RNA transcriptome in mutant SiHa C51S was first compared to SiHa. Most transcripts including E6, E7, and p53 were present at very similar levels in both cell lines. Interestingly, there were transcriptional differences with SiHa cells as depicted in the volcano plot in Supplementary Figure 4B. Together, these amount to less than 0.08% of all captured mRNAs. Following incubation of C51S cells in KTI-240, a total of 64 mRNAs increased and 109 decreased (Figure 5E; Supplementary Table 2). These 173 mRNAs may include changes resulting from the p53-independent activities of E6. In contrast to parental SiHa, incubation with KTI-240 did not demonstrate analogous changes in p53 responsive transcripts (Figure 5H). which underscores the compound’s reliance on Cys-51 for E6•E6AP mediated p53 degradation.

The RNA transcriptome changes in RPE-1 before and after incubation in KTI-240 were also characterized (Figure 5D, Supplementary Table 3). E6 and E7 mRNAs were not detected and low levels of p53 transcripts as in the SiHa cells were present. Only 25 genes were dysregulated following exposure to KTI-240 and six DEGs (SLC7A11, VSIR, NMRAL2P, OSGIN1, DHRS3, GCLM) overlapped with the changes in SiHa and C51S cells (Figure 6A). While these genes could be responding to KTI-240, none are associated with p53 induction. These results strongly infer that the cysteine reactive compound KTI-240 induced very limited HPV-independent effects.

**Figure 6.**
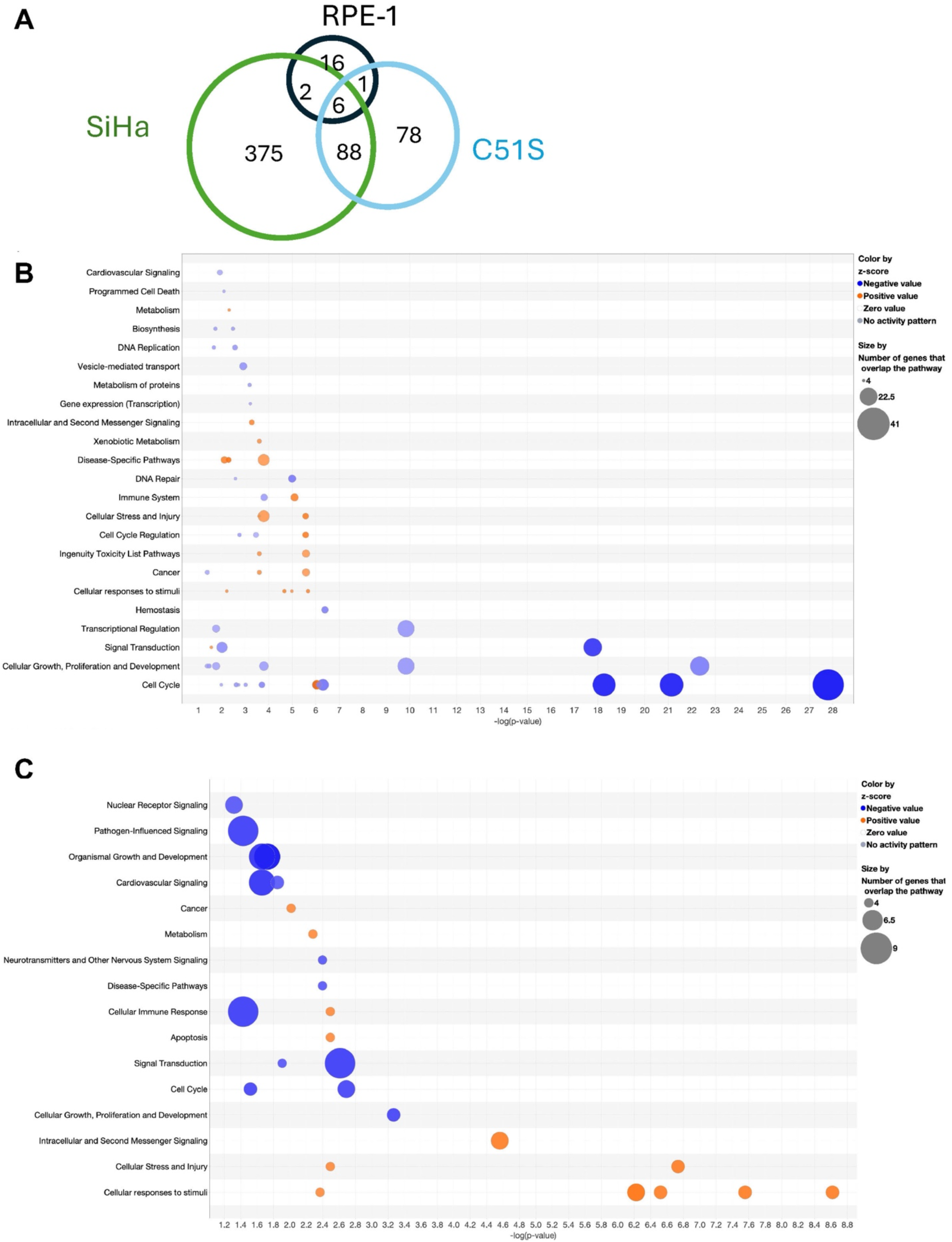
KTI-240 affects cell cycle and p53 pathways limited to wild-type E6 expressing cells. **(A)** Venn diagram of identified DEGs in SiHa, SiHa C51S, and RPE-1 cells. 25 DEGs were identified after KTI-240 treatment in RPE-1 cells, of which six overlapped with SiHa and C51S cells. SiHa had the most DEGs in response to KTI-240 treatment. **(B, C)** RNA seq data analyzed using Qiagen’s IPA pathway analysis. There were no significant pathways changed in RPE-1 cells, while SiHa cells displayed the most affected pathways. **B**) The bubble map shows that many affected pathways are cell cycle associated and are inhibited. This was not the case in C51S cells (**C**), where the majority were associated with cellular responses to stimuli and stress.

Signaling pathway analysis using Qiagen’s IPA software revealed reductions of cell cycle checkpoints, kinetochore metaphase signaling, and mitotic prometaphase and metaphase pathways in SiHa cells post KTI-240 treatment (Figures 6B, 7B). The identified network highlights the role of CDKN1A (p21) in inhibiting cell cycle progression by decreasing entry into interphase and reducing cell viability, leading to increased cell death and necrosis (Figure 7A). This interpretation aligns with the observed outcome when HPV16 E6 mediated p53 degradation is inhibited. Similarly, while TP53 transcript levels were unchanged, IPA analysis predicts a central role for p53 in this E6 inhibition network, influencing multiple pathways by increasing CDKN1A, IFNG, IL1A, IL1B, and TNF, while decreasing EP400 and RABL6. The observed increase in IL1A and IL1B in response to KTI-240 is consistent with the reported repression of these genes by HPV16 E6 and E7 during various stages of the HPV pathology including the late stages of carcinogenesis (32). Altered expression of these inflammatory cytokines, which can increase apoptosis and cell death in tumor cell lines, likely contributes to the regulation of apoptosis in SiHa cells. The network also shows connections between necrosis and apoptosis. Importantly, cell cycle and p53 pathways transcripts were unchanged in HPV-negative RPE-1 cells by KTI-240 (Figures 8A, B, C), and IPA pathway analysis did not detect a single significant pathway alteration. On the other hand, a significant number of pathways were affected in C51S cells after KTI-240 exposure. The cell cycle checkpoint pathway was similarly inhibited as in wild type SiHa cells (Figures 6C and 7D) albeit to a lesser degree, with fewer genes being affected (Figure 8C). Importantly, p53 signaling pathway transcripts were not dysregulated in KTI-240-treated C51S cells (Figure 8B), and neither p53 nor p21 were identified as major responders in the IPA analysis (Figure 7C). There were overlaps in a few pathways with low z-scores (Figure 8A). Overall, affected pathways involved fewer genes than in SiHa cells. We speculate that the transcriptional changes shared by wild-type and C51S SiHa cells are mediated by E6 interactions independent of the E6•E6AP•p53 complex. One such candidate is the transcription factor ATF3 (33).

**Figure 7.**
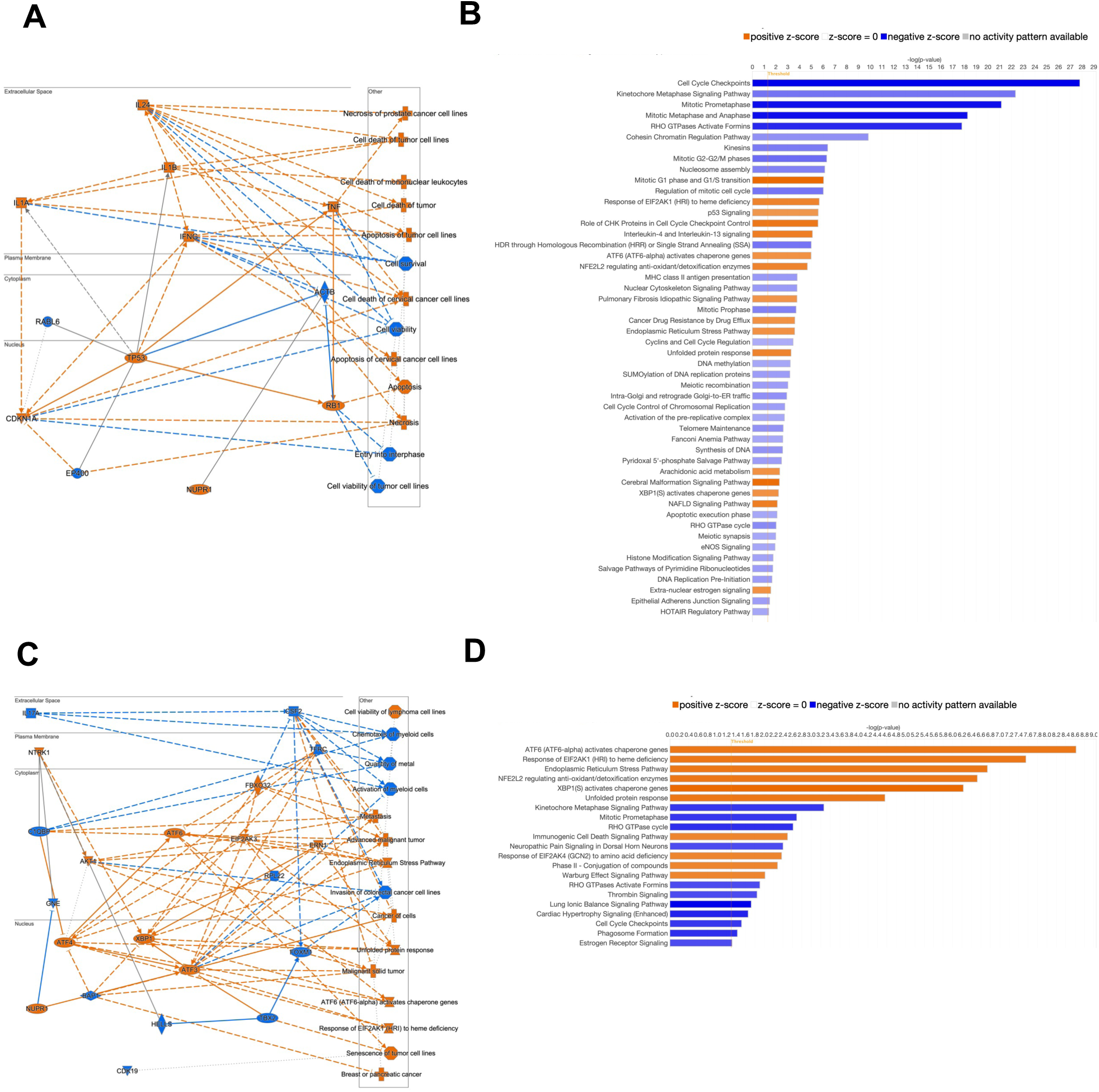
IPA pathway analysis identifies p53 as a key regulator of cell death associated pathways in KTI-240-treated SiHa cells. **(A)** Network analysis of differentially regulated pathways in SiHa cells. Importantly, TP53 is predicted to be upregulated and to be a key player of increased apoptosis, necrosis, and senescence pathways in KTI-240-treated SiHa cells. Individually significantly affected pathways are shown in **(B)**, where blue indicates inhibition and orange activation of the respective pathway. In contrast, p53 was not predicted to be a key player in KTI-240-treated C51S cells **(C, D)**. Overall, fewer pathways and DEGs were detected in treated C51S with ATF3 being a key player, which has been shown to directly bind to HPV16E6 and compete with p53 binding.

**Figure 8.**
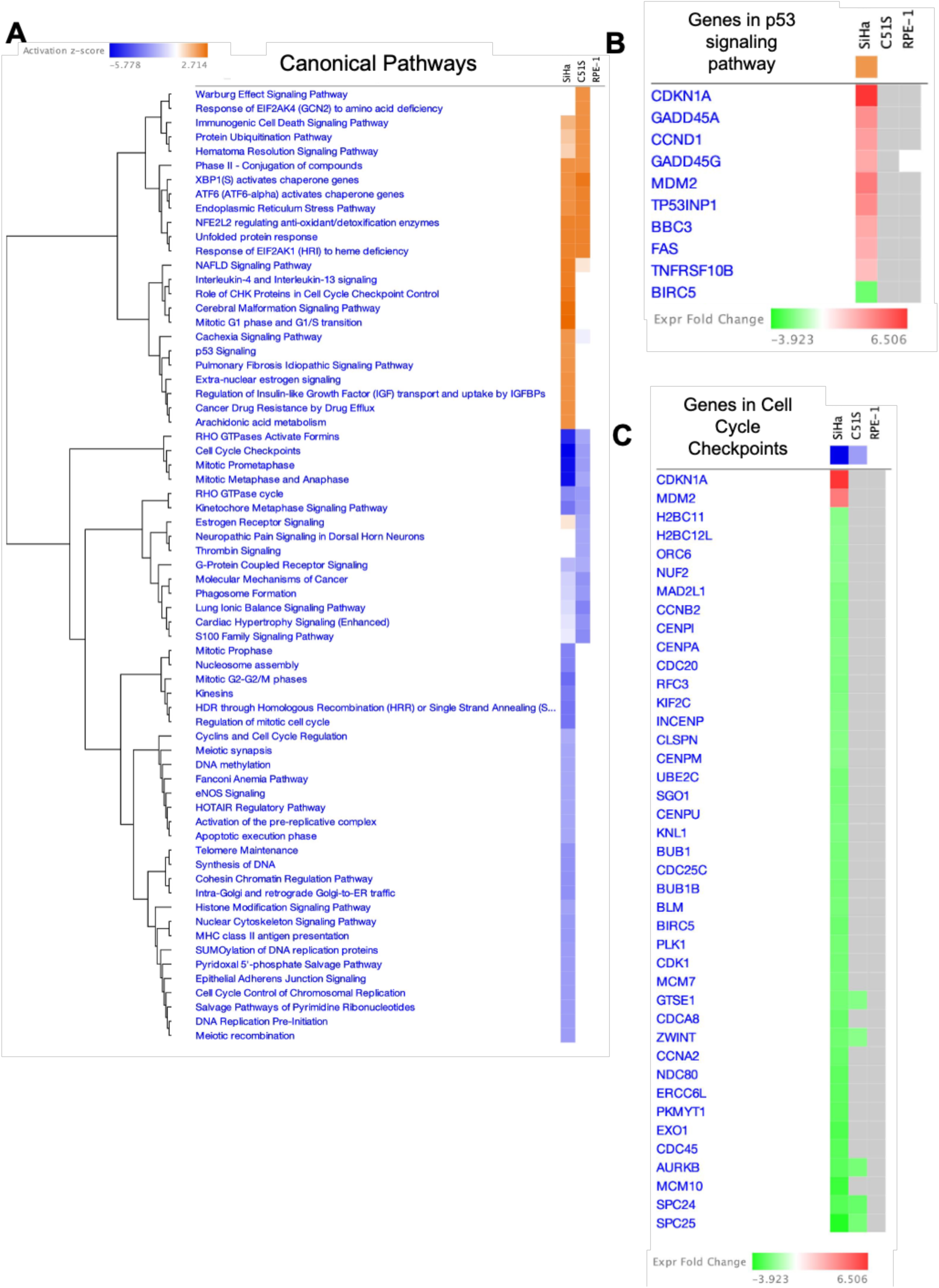
Comparison analysis of significantly changed pathways and genes in SiHa, C51S and RPE-1 cells. **A**) Comparison analysis showing a heatmap of differentially regulated canonical pathways between SiHa, C51S and RPE-1 cells, where orange indicates activation and blue inhibition. Heatmap of fold change gene expression of genes in the p53 **(B)** and cell cycle checkpoints **(C)** signaling pathway in SiHa, C51S and RPE-1 cells. Red indicates increased and green indicates decreased mRNA expression, grey indicated no significant difference was detected and white means mRNA was not detected or too low expressed.

We compared levels of representative proteins involved in HPV16 E6/E7 mediated carcinogenesis in SiHa, RPE-1, and C51S cells following 16 hours in KTI-240 supplemented media. CDKN1A (p21), which increased by ∼7-fold at the mRNA level (Figure 9E), showed a 3-fold elevation at the protein level in parental SiHa cells (Figure 9A, F; Supplementary Figure 5A) and were unchanged at the mRNA or protein level in RPE-1 after KTI-240 treatment (Figures 9A, E, F). We also confirmed both E6 binding compounds increase levels of p21 in HPV16+ OPC cells (Figures 4A, B). In C51S cells, KTI-240 did not increase p21 protein levels, although this mRNA was modestly elevated by 1.5 FC with an FDR of <0.05. In addition, p16 (CDKN2A) was unchanged at the mRNA or protein level. Similarly, E6AP mRNA and protein expression remained baseline after KTI-240 in both SiHa cell lines (Figures 9B, E, F). Activation of 53 stimulates MDM2 transcription which in turn induces p53 protein degradation in a negative feedback loop. Interestingly, while there was a 4-fold induction in MDM2 transcript levels in SiHa cells by KTI-240 (Figure 9E), this increase did not correspond with its protein levels. FOXM1 mRNA was significantly downregulated in SiHa and C51S cells with a corresponding decrease at the protein level by ∼50% but remained unchanged in RPE-1 cells (Figures 9E, C, F; Supplementary Figure 5E). Rb1 transcript levels were unaltered while Rb protein decreased in SiHa and C51S cells following KTI-240 exposure, but this did not occur in RPE-1 (Figures 9C, F). Both BBC3 (PUMA) mRNA and protein increased by 2.7 and 2.8-fold respectively in SiHa but were unchanged in RPE-1 and C51S cells (Figures 9D, E, F; Supplementary Figure 5G).

**Figure 9.**
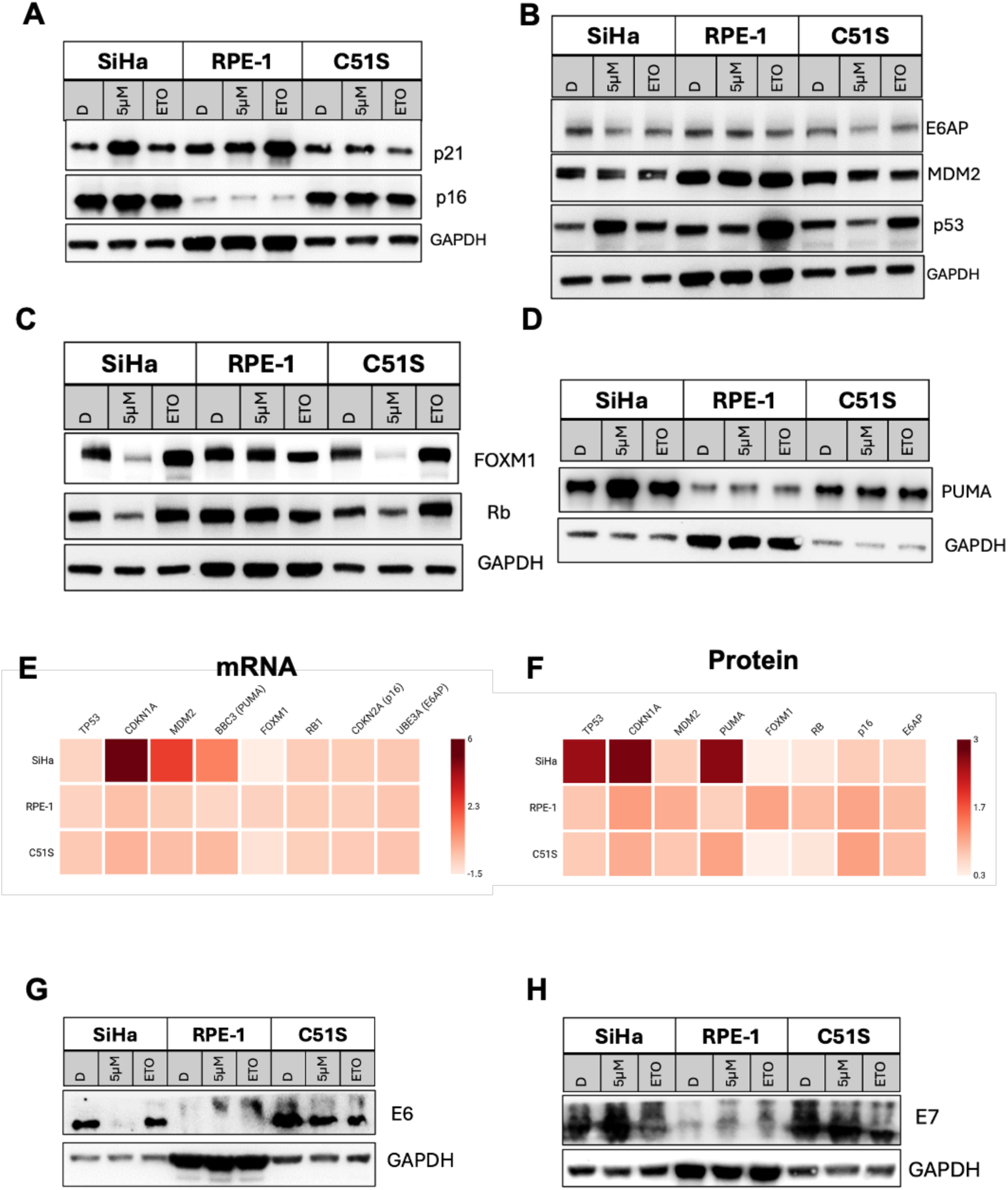
Validation of protein expression of key players in the HPV oncogenesis. SiHa, SiHa C51S, and RPE-1 cells were cultured in KTI-240 (5 µM), DMSO or etoposide (ETO, 25 µM) for 16 hours, cells were lysed, proteins extracted and subjected to Western blot analysis. **A)** KTI-240 increased p21 levels without affecting p16 protein. **(B)** E6AP and MDM2 protein levels were not changed after KTI-240 treatment. p53 protein was included as a positive control. KTI-240 decreased Foxm1 and Rb protein expression in SiHa and C51S cells but not RPE-1, while Puma increased only in SiHa cells. The heatmaps in **(E)** and **(F)** summarize the detected changes at the mRNA and protein level. **(G, H)** E6 protein levels were significantly decreased in SiHa cells, while E7 was unchanged in both HPV+ cell lines. Protein expression was normalized to GAPDH and expressed as fold-change. Data was expressed as S.E.M and analyzed using one-way ANOVA with Dunnett’s or Bonferroni post hoc analysis were applicable. Experiments were repeated at least three independent times (n ≥ 3; * p < 0.05).

The half-life of the E6 protein is dependent on association with E6AP (34). We questioned whether compound binding to E6 would alter its levels, and indeed Western blots showed a significant reduction in HPV16 E6 protein levels in SiHa cells but not in C51S cells after 16 hrs culture in KTI-240 (Figure 9G, Supplementary Figure 5H). While E6AP binds to and stabilizes the E7 protein (35), altered E7 protein levels were unaltered after KTI-240 exposure (Figure 9H; Supplementary Figure 5I).

### E6 binding compounds specifically reduce HPV cell viability

Two human cervical cancer cell lines, SiHa and CaSki (ATCC^®^ CRL-1550™), and two human cancer derived lines, UM-SCC-47 and UM-SCC-104, were cultured in increasing concentrations of KTI-218. These HPV16+ cells showed decreased viability with IC50 values between 1 and 3 µM after 24 hours of KTI-218 exposure, in contrast to primary human foreskin keratinocytes (HFK) and RPE-1, which showed a modest decrease in cell numbers at the highest concentrations of the E6 inhibitor (Figure 10A). Similarly, KTI-239, the non-covalent analog of KTI-218, did not decrease SiHa cell viability (Supplementary Figure 6A), supporting a requirement for covalent binding to E6 to drive cell death.

**Figure 10.**
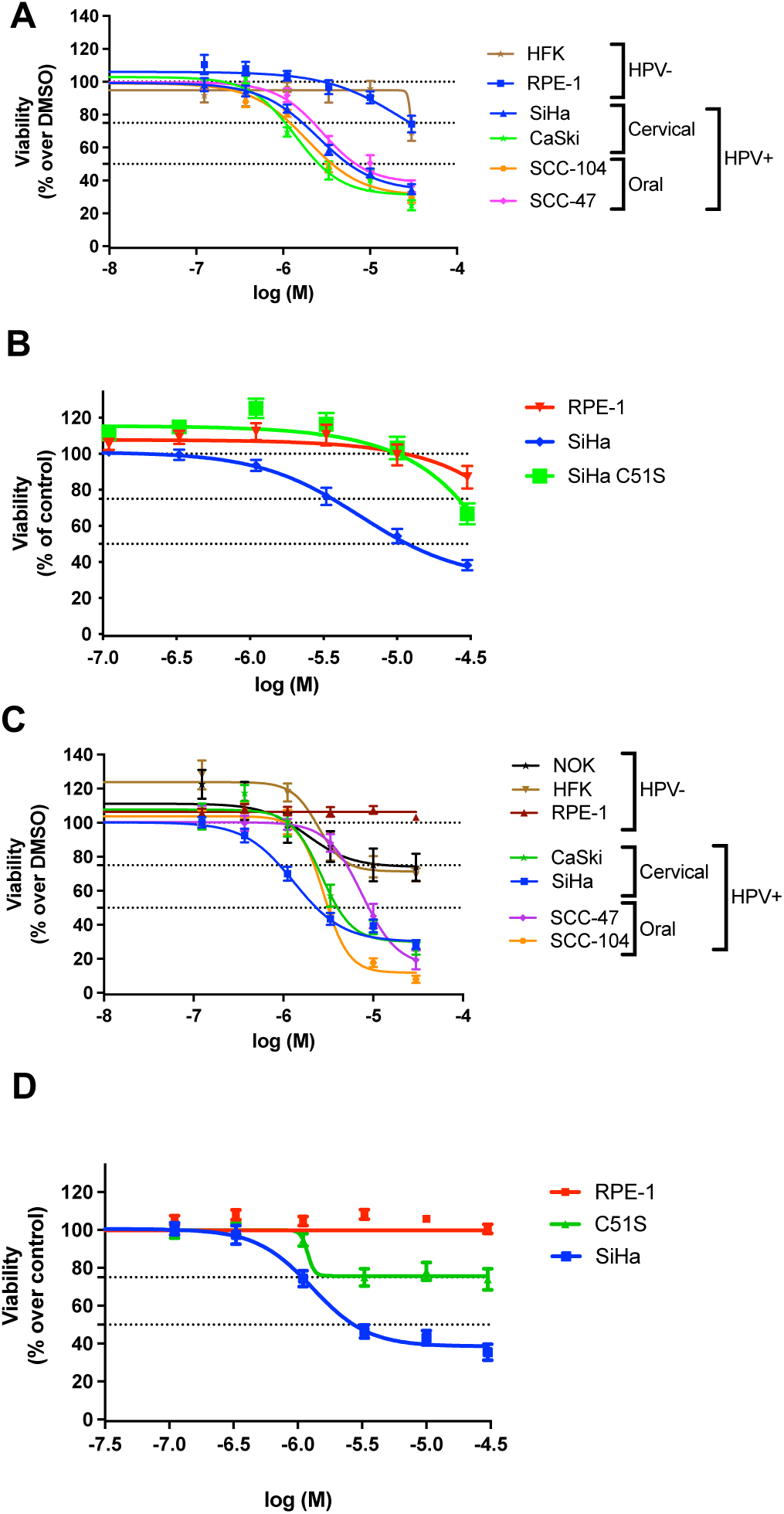
KTI-218 and KTI-240 specifically decrease viability of HPV+ cells. **(A)** HPV+ cervical cancer cell lines, SiHa and CaSki, the oral cancer derived lines UM-SCC-47 and UM-SCC-104, and RPE-1 were incubated with increasing concentrations of KTI-218 or DMSO for 24 hours and cell viability was analyzed by Calcein-AM assay. **(B)** HPV negative RPE-1, human foreskin keratinocytes (HFK) and C51S mutant cells were analyzed for their cell viability after KTI-218 treatment. **(C, D)** cell viability of SiHa, CaSki, SCC-47, SCC-104, SiHa C51S, and RPE-1, HFK and normal oral keratinocytes was assessed after KTI-240 or DMSO treatment with for 48 hours. Cell viability is expressed as a percent change over DMSO control-treated cells. Data expressed as S.E.M and each experiment was completed at least three independent times (n ≥ 3).

After cells were exposed to KTI-240 for 24 hours, a significant but more modest reduction in viability was observed in SiHa cells (Supplementary Figure 6B), while RPE-1 cells were unaffected. This series of HPV+ and HPV-negative cells were exposed for 48 hours to KTI-240, which decreased cell viability in cervical and oral HPV+ cells with IC50s between 1 and 3 µM for SiHa, CaSki and SCC-104, while SCC-47 were less sensitive to KTI-240 with an IC50 of 7.4 µM. KTI-240 did not decrease viability of RPE-1 cells and caused slightly decreased viability of HFKs and normal oral keratinocytes (NOK) at the highest concentrations (Figure 10C). Given the high induction of p53 in W12-E cells after KTI-218 and KTI-240 treatment, we found a concordant significant reduction in cell viability (Supplementary Figure 6C), implying that inhibition of HPV16 E6 in cells with episomal HPV genomes causes cell death. The transcriptomic data predicted the inhibition of some cell cycle death pathways in C51S cells albeit to a lesser extent than in SiHa cells in response to KTI-240. Indeed, there was a modest reduction in C51S cell viability in KTI-240 or KTI-218 supplemented media, but to a much lesser extent than the parental SiHa cells (Figures 10 C, D).

The RNA-seq data documented activation of a cell death pathway that involves apoptosis in SiHa cells. KTI-240 specifically increased cleaved caspase 3 by 3.5-fold in SiHa cells but not RPE-1 cells, consistent with cell viability studies (Figures 11A, B). Interestingly, C51S SiHa showed a smaller 1.6-fold increase in apoptotic cells. To further investigate this finding, we measured apoptosis in cells in the absence of drug. Interestingly, there was an ∼15% rate of basal apoptosis in C51S cells compared to 5% in parental SiHa cells (Figure 11C; Supplementary Figure 7). KTI-240 (3, 10 µM) treatment for 48 hrs significantly increased apoptosis in SiHa cells but only modestly in C51S cells (Figure 11D). Next, we investigated if KTI-240 induces senescence in these three cell lines. Basal β-galactosidase staining was negligible in SiHa and RPE-1 cells, while SiHa C51S cells had significantly higher levels of basal β-gal staining intensity compared to parental SiHa cells (Supplementary Figure 7B). Expression of SA-β-gal was induced in etoposide treated SiHa and C51S cell lines. KTI-240 increased this senescence marker in SiHa cells but not in RPE-1 cells and only modestly in C51S (Figure 11E; Supplementary Figure 7B, C).

**Figure 11.**
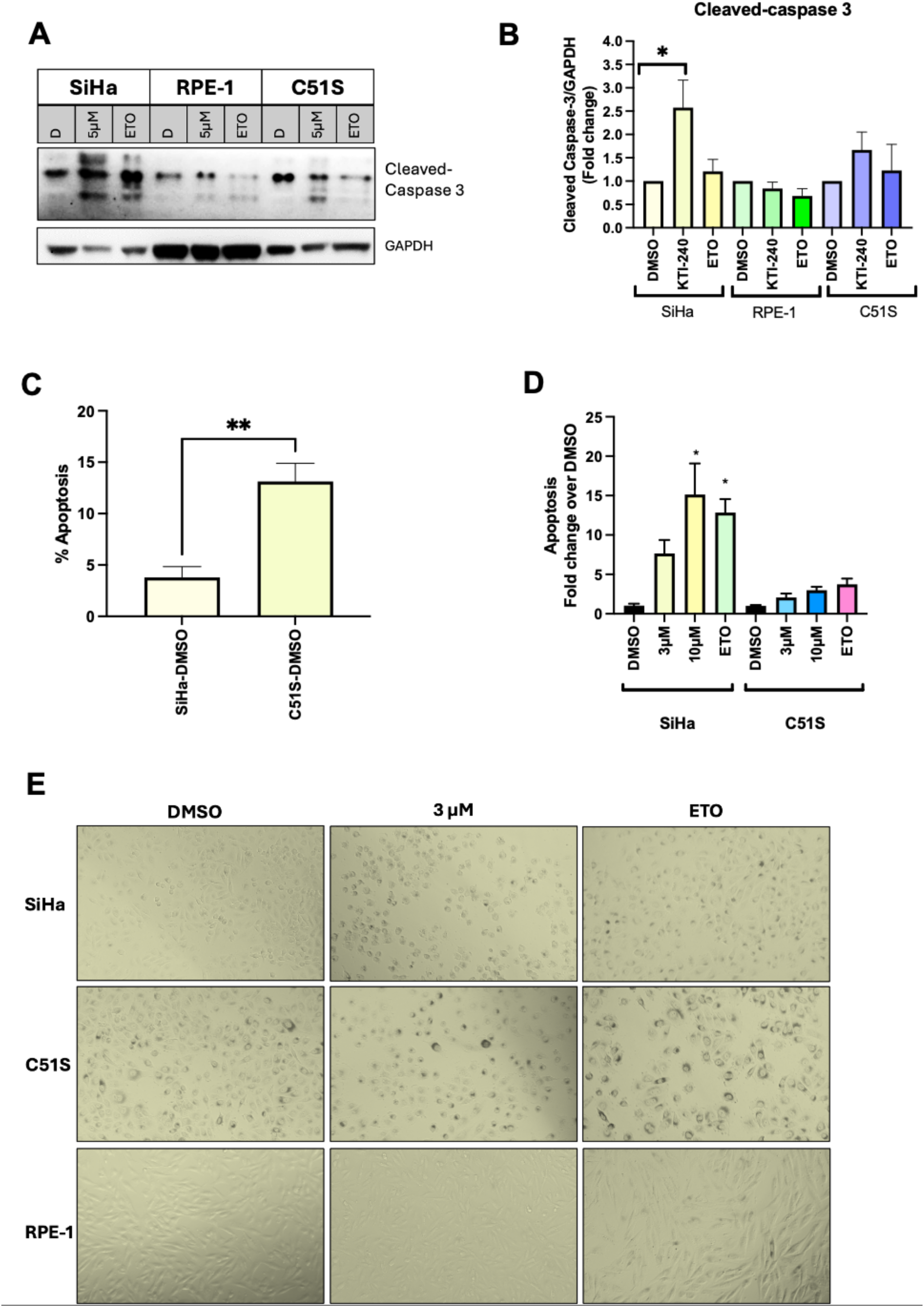
KTI-240 specifically increases apoptosis and senescence in HPV+ SiHa cells. (**A, B**) HPV+ SiHa cells, C51S mutant and HPV-negative RPE-1 cells were incubated with DMSO, 5 µM KTI-240 or 25 µM etoposide for 48 hours. Cells were lysed and extracted proteins were subjected to Western blot analysis for cleaved Caspase 3 protein. (**C, D**) Cells were treated for 48 hours with DMSO, KTI-240 (3 and 10 µM) or etoposide (2 µM) and apoptosis quantified with FITC-Annexin V. **(E)** Senescence was analyzed by a SA-β-gal assay after 48 hours of KTI-240 (3 µM), etoposide (2 µM) or DMSO treatment. Protein expression was normalized to GAPDH and expressed as fold-change. Annexin-FITC and β-Gal staining intensities were normalized to DMSO control. Data is expressed as S.E.M and was analyzed with one-way ANOVA with Dunnett’s or Bonferroni post hoc analysis or unpaired Students t-test (**C**). Each experiment was completed at least three independent times (n ≥ 3; * p < 0.05).

### In Vivo Tumor Xenotransplantation experiments

HPVs are species-specific and do not infect other animals including rodents. We decided to use authentic human sourced cancer cell lines that express E6 and E7 from the native HPV promoter and which are known to be dependent on maintaining these viral proteins (36). Prior to this series of ‘proof of concept’ experiments, *in vivo* bioavailability of KTI-218 after intraperitoneal injection (IP) into C57BL/6 mice was measured. KTI-218 was administered at a dose of 50 mg/kg and plasma levels were measured over 24 hours (Supplementary Figures 8A, B). KTI-218 showed an acceptable pharmacokinetic profile to undertake key proof of concept *in vivo* studies. Initial testing was performed in a xenograft model with the oral HPV+ cancer cell line UM-SCC-47. Treatment of male Nu/Nu mice with 50 mg/kg KTI-218 or vehicle per day began ∼ 7 days post-implantation when the tumors were 50-100 mm^3^. Mice were treated once daily, 6 days a week.

Administration of KTI-218 significantly slowed tumor growth without affecting body weight (Figure 12A, B; Supplementary Figure 8C). To visualize the viable tumor cells, we injected a subclone of the cervical HPV+ SiHa cell line that constitutively expresses luciferase (Luc) and which exhibits similar growth properties as parental cells. Luciferase expression reflects the presence of viable tumor cells and is quantified by detection *in vivo* using the IVIS SpectrumCT scanner that can be performed on live mice. This experiment demonstrated that some mice did not exhibit measurable luciferase activity at day 39, while others showed significantly less activity than vehicle treated animals (Figures 12C, D). Overall, the measured Luc activity correlated well with the caliper-measured tumor sizes. Body weight was not impacted by the administration of KTI-218 (Supplementary Figure 8F). When animals reached their endpoints, mice were sacrificed, and the remaining nodules were extracted and weighed (Figure 12E). Tumors of drug-treated animals were significantly smaller and weighed less than the vehicle-treated tumors. To test if the remaining viable cells developed resistance towards the drug, we isolated cells from two tumors and propagated these in culture. These cells were treated with KTI-218 and found to be equally sensitive as the injected SiHa cells (data not shown). These results infer that persistence of tumor cells is not due to development of drug resistance. In parallel, we also isolated DNA and sequenced PCR-amplified DNA from a small residual UM-SCC-47 tumor and found that the cysteine 51 coding sequence for HPV16 E6 was preserved. We also tested the efficacy of KTI-218 in male Nu/Nu mice xenotransplanted with SiHa cells. Here, mice were treated with 50 mg/kg KTI-218 or vehicle per day and began ∼14 days post-implantation when the tumors were 50-100 mm^3^. KTI-218 also significantly slowed tumor growth in this model (Supplementary Figure 8D, E). In contrast, KTI-218 did not affect the tumor growth of HPV negative cervical cancer C33a cells transplanted into female nude mice (Figure 12F). Next, we tested the efficacy of KTI-240 on tumor growth of SCC-47 xenografts. When administered once daily at 25 mg/kg, inhibition of SCC-47 tumor growth was not observed (data not shown). However, by increasing the dosage of KTI-240 to 50 mg/kg/day, reduced tumor growth of SCC-47 xenografts was achieved (Figure 12H). Two hours after the final dose, mice were sacrificed and tumors analyzed for drug levels. KTI-240 was detectable in the tumor tissue with an average concentration of 1.5 µg/gm. Importantly, we inoculated nude mice with SiHa C51S cells, which formed tumors with similar growth as wild-type SiHa. The C51S tumors enlarged comparably in mice treated with KTI-218 by daily IP injection or with vehicle alone (Figure 12G). DNA sequencing confirmed the presence of the serine codon in the tumor tissue. These results further evidence the importance for covalent bond formation to Cys51 in mediating inhibition of E6 activity *in vivo*.

**Figure 12.**
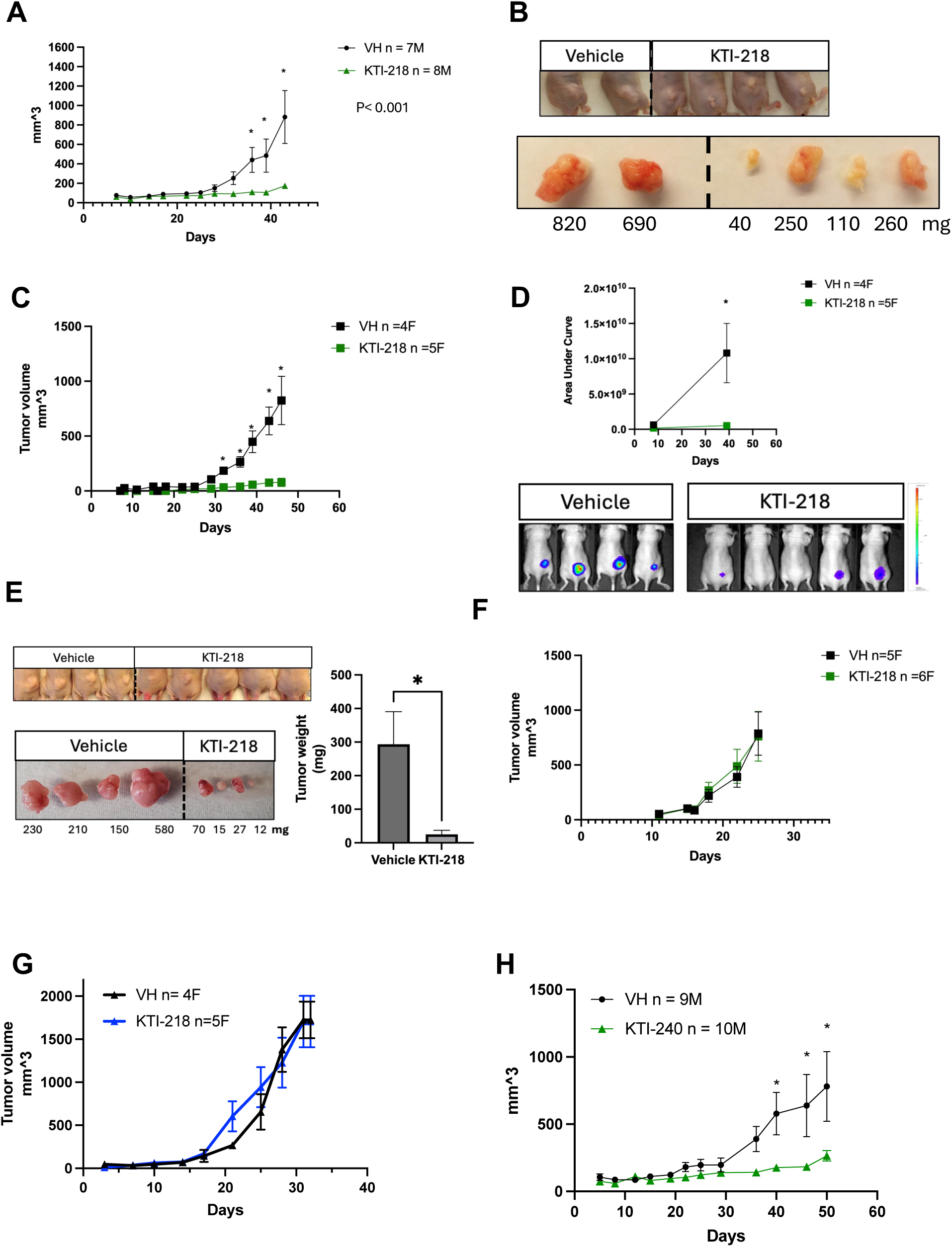
KTI-218 and KTI-240 inhibit growth of HPV-16 expressing human cervical and oropharyngeal tumor cells *in vivo*. (**A, B**) Nu/Nu mice were injected subcutaneously with HPV-16+ SCC-UM-47 cancer cells. Intraperitoneal injections of 50 mg/kg KTI-218 (n=8, green) or vehicle (VH, n=7 black) began when tumors were ∼50-100 mm^3^. Tumor size was measured by calipers and expressed as S.E.M. and analyzed using two-way ANOVA (p<0.01). Representative mice and extracted tumors are shown in (**B**). Nu/Nu mice were injected subcutaneously with HPV-16+ SiHa cancer cells stably expressing a Luciferase construct (**C**). Intraperitoneal (IP) injections of 50 mg/kg KTI-218 (n=5, green) or vehicle (VH, n=4 black) began when all tumors had a bioluminescent signal. Bioluminescence was measured using a IVIS SpectrumCT optical imaging system (**D**). Representative tumors and tumor weights are shown in (**E**). Nu/Nu mice were injected subcutaneously (SC) with (**F**) HPV-negative cervical cancer cell line C33a or (**G**) HPV + SiHa C51S cells. Intraperitoneal injections of 50 mg/kg KTI-218 began when tumors were ∼50-100 mm^3^. (**H**) Nu/Nu mice were injected SC with HPV-16+ SCC-UM-47 cancer cells. IP injections of 50 mg/kg KTI-240 (n=10, green) or vehicle (VH, n=9 black) began when tumors were ∼50-100 mm^3^. Data expressed as S.E.M and analyzed using two-way ANOVA; * p <0.05.

## DISCUSSION

The E6 protein is essential for maintenance of HPV episomal replication in epithelial cells and along with E7 is always expressed in HPV associated malignancies. Reductions in E6 levels by siRNA or CRISPR targeting in tumor cells have been reported to induce apoptosis and senescence (37, 38). When expressed alone, HR E7 increases and activates p53 (39–41). Therefore, E6 is an attractive target for antiviral therapy to specifically eliminate HPV infected cells and avoid the possibility of developing resistant mutations (42). Prior attempts have not successfully identified drug-like molecules that block E6 functions *in vivo* (13, 43–48). For the present strategy, we noted a cysteine residue proximal the binding pocket of HPV16 E6 for alpha-helical LXXLL motif proteins, which offered the potential to develop compounds with a warhead that covalently bonds to this residue and irreversibly blocks E6 binding to its cellular partners. Although most if not all HR and many other HPVs bind to E6AP, only high-risk HPV-35, 45 and 68 encode an analogously positioned cysteine. While variant amino acids in HPV16 isotypes have been detected in benign and malignant lesions (49), these are uncommon and mostly uncharacterized. To the best of our knowledge, variants of cysteine 51 in HPV16 E6 have not been observed.

The HECT domain ubiquitin ligase E6AP associates with E6 via a helical LXXLL motif. Other regions of E6AP contact E6 and contribute to overall binding affinity (50). Differing *in vitro* binding affinities have been reported and may be in the picomolar range (51). Cryo-EM structures of HPV16 E6 in complex with the full E6AP and partial p53 proteins indicate that relatively little of the E6 surface is exposed (52–55). Whether the LXXLL binding domain of E6 would be accessible by a small molecule and could compete with the high affinity E6AP protein was questioned. To circumvent this consideration, we designed molecules with a warhead to covalently bond to cysteine 51. Another group has inhibited E6 binding to E6AP using a peptide modified to covalently bond to C51 (called C58 in this publication) (51). The results we present herein demonstrate successful inhibition of E6 can occur in live cells and *in vivo*. One explanation is highly purified E6 and E6AP proteins produced *in vitro* do not recapitulate their biological interactions in living cells. Both E6 and E6AP have been shown to be post-translationally modified. Another potential mechanism of inhibition is that freshly translated E6 protein may bind and rapidly covalently react with the compounds before encountering E6AP. Alternatively, compound binding may impede the LXXLX binding interaction and yet E6AP may still complex with E6 due to the additional sites of protein-protein interaction, however this conformation may not be competent to mediate p53 ubiquitination. It is also possible that the E6 inhibitory compounds interfere with a conformational shift in E6 necessary for binding to p53 or may directly block p53 association or its ubiquitination. Future experiments will delineate these mechanistic possibilities. Nonetheless, our accumulated data demonstrate that KTI-218 and KTI-240, each containing a reactive acrylamide warhead that bonds to a specific cysteine in E6, effectively and selectively increase p53 levels and activities in E6 expressing cells.

E6AP association was shown to increase the half-life of the E6 protein (34) and E6AP was also reported to bind to HPV E7 protein and regulate its stability (35). In our experiments, KTI binding to E6 significantly reduced E6 protein levels without affecting E6 or E7 mRNA levels or E6AP mRNA and protein levels. Therefore, we speculate that covalently bound KTI compounds destabilize the E6 protein by interfering with E6AP association. An atomic structure of E6 in complex with a small molecule has not been previously reported. Co-crystallization experiments with covalently bound compounds are currently underway and may illuminate the causative mechanism.

E6 inhibition led to robust elevations in p53 protein levels and induction of p53 transcriptional activities. The decreased cell viability limited to HPV16 E6 expressing cells exposed to these compounds can be partially explained by the restoration of p53. P53 transcriptional activities are regulated by an assortment of post-translational modifications. We did not attempt to discern which of these might be occurring in the KTI treated HPV16 cells. In experiments using siRNA protocols, the interaction of E6 with E6AP was reported to be necessary for blocking cell senescence and was attributed to increased expression of p21 (16). Our chemical inhibitors of E6 are likely to be more effective due to the combined induction of p53 pathways of apoptosis and senescence.

Remarkably few transcriptome alterations were identified in telomerase immortalized RPE-1 cells that express wild-type p53 following culture in KTI-240. The CRISPR-edited SiHa cells with the cysteine 51 mutated to-serine (C51S) did not manifest downstream transcriptional consequences of p53 activation and were only slightly susceptible to growth inhibition by these compounds, despite that these maintain the E6•E6AP mediated p53 degradation pathway and are tumorigenic. Their slight decrease in proliferation and viability in response to KTI-218 and KTI-240 may be due to the p53-independent effects of E6 mediated through its LXXLX binding motif or other interaction partners that contribute to cancer cell proliferation (56, 57). Structural modeling suggests it is unlikely that these compounds obstruct the E6 C-terminal PDZ binding motif (58, 59). We also noticed that the basal transcriptome of the C51S cells had differences from the parental SiHa. Our interpretation is that the C51S line expanded from a single cell and that commercially available SiHa cells are not a completely homogeneous population. We also compared our RNA seq data to a published study [accession # GSE235662 in (60)] in which HPV16 E6 and HPV16 E6+E7 were transduced into ‘head and neck cells (HNC)’ with the idea that pathways affected by E6 inhibition would be oppositely influenced by E6 or E6/E7 overexpression. Indeed, there was a good correlation with an inverted profile of affected signaling pathways (Supplementary Figure 9).

We chose to use authentic human HPV-cancer derived cell lines that depend on E6 and E7 for *in vivo* xenotransplantation experiments. This required using immunodeficient murine hosts, which limits potential immune responses following release of viral antigens from dying cells. E6 binds to IRF-3 (61) and interferes with cell autonomous immune recognition (62), which may be important in human host responses but not available in the nude mouse. While we considered the inducible LTR driven HPV16 E6+E7 transgenic OPC mouse models (63–65), tumors appear several months following Cre-induced expression and require the concomitant expression of mutant Ras or PIK3CA. Furthermore, the interactions of HPV E6 with mouse proteins including E6AP and p53 may not faithfully mirror the dynamic structural changes that have been reported using purified proteins. Dramatic reduction in tumor growth was restricted to wild-type E6 expressing cells and was not observed with the C51S SiHa derivative. The persistence of small residual tumors is suspected to be result of inadequate vascularity and drug availability. Isolates of these remaining cells showed retention of susceptibility to drug and absence of mutation in the E6 gene.

The HPV16 genotype is the prototypic high-risk (HR) PV and accounts for nearly 50% of all cervical cancers across the world. HPV16 is also identified in the majority of anal and genital intraepithelial dysplasias and cancers (66). Worldwide ∼100,000 oropharyngeal cancers (OPC) are attributable to HPV16 and in the USA HPV-positive OPC (>20,000/year) exceeds the incidence of cervical cancer. While the current HPV vaccine is >95% effective in preventing infection, it does not alter the prognosis or course of people already infected with the virus. Even though it became available in 2006, there has been insufficient uptake to achieve herd immunity in the USA and the vaccine is too expensive for widespread use in many countries where cervical cancer is the predominant cause of death of women. It has been predicted that 44 million cases of cervical cancer will arise over the next 50 years (67), highlighting the immense need for an antiviral remedy. We propose that an E6 inhibitory drug could be useful in multiple clinical scenarios including cervical, genital, and anal cancers that have poor outcomes after metastatic spread. While treatment of HPV16 association OPCs is associated with a relatively high five-year survival, the morbidity of the current surgical and radiotherapy protocols is also extremely high. Because E6 inhibition results in high levels of wild-type p53 protein, this may increase susceptibility of HPV16 malignancies in combination with standard protocol DNA damage therapeutics such as x-ray and chemotherapy. Robust induction of p53 in the human cervical dysplasia derived cell line W12-E suggests that E6 inhibition may also be a viable strategy for premalignant pathologies. It is interesting to note that retention in the basal epithelium is attributable to E6 expression in keratinocytes (68, 69), and abrogation of this function would interfere with viral persistence. Cervical dysplasia afflicts millions of women annually and elimination of these HPV16 infected tissues may offer a novel non-surgical approach to prevent cancer in infected hosts.

## LIMITATIONS OF THE STUDY

Use of established human cancer derived cell lines, which are known to be dependent on E6 and E7 expression, may have accumulated other mutations that might affect drug responsiveness. The antitumor effects reported here might not be representative of naturally arising HPV cancers. Dose ranging studies were not performed.

## RESOURCE AVAILABILITY

Further information and requests for resources and reagents should be directed to and will be fulfilled by the lead contacts, Anne Rietz and Elliot Androphy.

## MATERIALS AVAILABILITY

All unique/stable reagents generated in this study will be made available on request with a completed materials transfer agreement.

## ACKNOWLEDGEMENTS

This work was supported by R01CA252715, subcontracts from R44CA268137, R43DE031495, and R43AI167573, the Mary Kay Ash Foundation, the Indiana CTSI 5UM1TR004402, and Kovina Therapeutics Inc. We appreciative work performed by the IU Genetics Core to generate the C51S cell line, the IU CMG and CCBB cores for RNA sequencing, IU CPAC core for determination of plasma drug levels, and the support of IU Simon Cancer Center’s CD3a program. NOK, human keratinocytes, and W12-E cells were generously shared by Karl Munger, Rachel Katzenellenbogen and Paul Lambert, respectively. We thank Thomas Raub, Essa Siddiqui, Steven Brooks, Susanna Tsueda, and Jacob Astroski for their dedicated technical assistance. The content of this manuscript is solely the responsibility of the authors and does not represent the official views of the Indiana University School of Medicine or the NIH.

## AUTHOR CONTRIBUTIONS

A.R. and EJA designed the experiments and interpreted data. A.R. and L.K. performed the experiments. A.K. and S.P. generated the C51S clone, Z.L. implemented the SAR studies that identified KTI-218 and KTI-240, which were supplied by Kovina Therapeutics Inc. A.R and E.J.A. drafted the manuscript.

## METHODS

### Cell Culture

SiHa, C51S, CaSki, UM-SCC-47, UM-SCC-104), C33a and RPE-1 cells were cultured in Dulbecco Modified Eagle Medium (DMEM) medium, supplemented with 10% fetal bovine serum (FBS, Perk Serum) and 1% penicillin-streptomycin (Gibco). N-TERT normal oral keratinocytes (NOKs) and human foreskin keratinocytes (HFK) were maintained in Keratinocyte-SFM (Gibco) supplemented with human recombinant epidermal growth factor (rEGF) and bovine pituitary extract (BPE) with 1% penicillin-streptomycin. W12-E that maintain HPV-16 episomes were grown in F-medium with J2-3T3 fibroblast feeders. Cells were tested regularly for mycoplasma and confirmed to be uninfected. Cells were cultured in humidified 95% air and 5% CO_2_ at 37 °C. Cells were passaged every three days at ∼70-80% confluence by washing with 1X PBS followed by treatment with 0.05% trypsin/EDTA, except for UM-SCC-47 and UM-SCC-104 cells, for which 0.25% trypsin/EDTA was used. For NOKs and HFK cells, trypsin was inactivated with trypsin inhibitor or media containing 10% FBS.

### Generation of HPV16 E6 C51S missense mutation in SiHa cells

SiHa cells (ATCC, HTB-35) were engineered using CRISPR-Cas9 methodology (70). Guide sequence and homology directed repair template were designed as described (71). SiHa cells were electroporated using the Amaxa 4D Nucleofector (Lonza) with the SE buffer and EN-138 program with a plasmid encoding Cas9-P2A-GFP and a single guide RNA targeting the E6 gene of the HPV16 (HPV16_E6_C51S_G1) together with a protected homology-directed repair (HDR) template carrying the C51S missense mutation flanked by short homology arms. Forty-eight hours post-transfection, GFP-positive cells were single-cell sorted into 96-well plates. Clones were expanded for 14–21 days, then replica-plated for genomic DNA extraction, genotyping, expansion, and cryopreservation. Correctly targeted clones were identified by PCR amplification and Sanger sequencing.

### Luciferase and viability multiplex assay

SiHa and RPE-1 cells were transfected with p53-renilla-luciferase (pRluc-C2; Perkin-Elmer) using polyethylenimine (PEI). Stable SiHa (denoted as SiHa p53-Luc) and RPE-1 (denoted RPE-1 p53-Luc) luciferase cell lines were established by 1.5 mg/ml geneticin selection. SiHa p53-Luc and RPE-1 p53-Luc cells were seeded in complete DMEM at 7,000 or 4,000 cells per well in white plates for 24 or 48 hrs drug exposure studies, respectively. The following day, media was replaced to DMEM with 1% FBS containing varying concentrations of KTI compounds or DMSO. DMSO concentration was kept constant at 0.02% (v/v). After 24 or 48 hrs, media was removed and 50 µl of PBS containing 2 µM Calcein-AM (calcein-acetoxymethyl ester) was added per well and incubated for 30 min at 37 °C. Calcein-AM is a cell-permeable fluorescent dye that is converted into green fluorescent calcein by intracellular esterases in viable cells. This conversion traps the dye within cells that have intact membranes, allowing for selective detection of live cells. Cell viability was determined by measuring fluorescence at an excitation/emission wavelength of 488 nm/520 nm. Following fluorescence measurements, PBS containing Calcein-AM was removed and renilla luciferase activities were measured using Dual-Glo® Luciferase Assay System (Promega, #E2920) according to manufacturer’s instructions. Renilla luciferase activity was normalized to cell viability and data expressed as percentage change over DMSO control. Assays were performed at least in triplicate and repeated at least three independent times.

### Cell viability assays

SiHa, RPE-1 and SiHa C51S mutant cells were seeded at 7,000 or 5,000 cells per well in black, clear-bottom 96-well plates for 24 and 48 hour drug exposure experiments. For all other cells, 10,000 and 7,000 cells per well were plated for 24 hour and 48 hour, respectively. Prior to drug treatment, medium was replaced with complete DMEM with 1% FBS, followed by the indicated concentrations of KTI-218 or KTI-240. Studies in HFKs, NOKs, and W12-E cells were performed in Keratinocyte-SFM supplemented with 1% FBS. Following incubation at 37 °C for 24 or 48 hours, the medium was gently removed and 50 µL of 2 µM Calcein-AM in PBS was added. Plates were incubated for 30 minutes at 37 °C and fluorescence measured using a microplate reader at an excitation/emission of 488/520 nm.

### Western Blots

20,000 or 15,000 cells/well for 24 and 48-hour experiments were seeded in 12-well plates. The following day, media was replaced to complete DMEM with 1% FBS containing varying concentrations of KTI compounds or DMSO (0.02% (v/v)). Cells were harvested and proteins were extracted in lysis buffer (10 mM Tris-HCl [pH 8], 150 mM NaCl, 2% SDS) supplemented with freshly added 1X protease inhibitor cocktail (SigmaFast, EDTA free). Protein concentration was determined with BCA Protein Assay Kit (Thermo Scientific). For most blots, 20 µg of protein lysate per lane was loaded, whereas 40 µg was loaded for cleaved caspase-3, 100 µg for HPV16 E6, and 80 µg for HPV16 E7. Samples were resolved on 4–12% or 4–20% SDS-PAGE gels using 1× MOPS running buffer and transferred to membranes using a semi-dry transfer system. Membranes were incubated for 45 minutes in 5% non-fat dry milk in Tris buffered saline (TBS)-Tween (0.1%) (TBST). Primary antibodies were diluted in 2% BSA/TBST and 0.02% NaAzide. Blots were incubated overnight, washed 3 times with TBST, and secondary antibodies added. HRP-conjugated secondary antibodies used in this study were: Peroxidase AffiniPure® Donkey Anti-Rabbit IgG (H+L), Peroxidase AffiniPure® Goat Anti-Mouse IgG, light chain specific and Peroxidase AffiniPure®, Donkey Anti-Mouse IgG (H+L) with dilution 1:2000. Detection was carried out using the SuperSignal™ West Femto chemiluminescence substrate for low expressed proteins and a conventional ECL detection system (GE Healthcare) for GAPDH and other higher expressed proteins. Chemiluminescent signals were acquired using a ChemiDoc Imaging System (Bio-Rad). Band intensities were quantified using ImageJ software. Experiments were repeated at least three times. GAPDH was used as loading control.

### siRNA-mediated knockdown of E6AP and E6/E7

Stable SiHa and RPE-1 luciferase cells were seeded as described in white 96-well plates, followed by transfection with 10 nM siRNA (control, E6-1, E6-2, E6AP, and E6/E7) and RNAiMax. Luc activities were measured 48 hr and 72 hrs after transfection. Parental SiHa and C51S cells were seeded at a density of 1.25×10⁵ cells/well in 12-well plates containing complete DMEM supplemented with 10% FBS. After 24 hrs, cells were transfected with siRNA targeting E6AP (UBE3A SilencerSelect siRNA, ID: s14604; Ambion,) HPV16 E6E7 (72), or a scrambled siRNA (SilencerSelect Negative Control # 1 siRNA; Ambion) at a final concentration of 15 nM using Lipofectamine RNAiMax (Life Sciences Technologies). After 72 hrs, cells were harvested and lysed in lysis buffer.

### DNA isolation and PCR amplification

Genomic DNA was isolated from cell pellets and mouse tumors using the DNeasy Blood and Tissue Kit (Qiagen). DNA concentration and purity were determined using a Nanodrop spectrophotometer. PCR was performed using HPV16 E6-specific primers (HPV16_E6_C51S_F51; HPV16_E6_C51S_R52) and RedTaq DNA Polymerase (Bullseye, BE180303) with the following conditions: 98 °C for 2 min; 35 cycles of 98 °C for 10 sec, 55 °C for 30 sec, 72 °C for 1 min; and a final extension at 72 °C for 5 min. Amplified PCR products were submitted for sequencing to ACGT, Inc. to confirm the presence of a cysteine codon at position 51 in 16E6.

### Senescence and Apoptosis Assays

40,000 cells per well were seeded in 24-well plates. The following day, cells were treated with varying concentrations of KTI-240 and 2 µM etoposide (ETO) for 48 hours in DMEM supplemented with 1% FBS. Vehicle-treated cells (0.02% DMSO) were included as control. For senescence analysis, the Cell Senescence β-Galactosidase Staining Kit (APExBIO). Cells were fixed, incubated with β-Gal staining solution at 37 °C without CO₂ overnight followed by imaging with a OLYMPUS IX73 optical microscope at a 20x magnification. For analysis of apoptosis, the Biotiums Apoptosis Kit was used. Cells were stained and imaged at 20x magnification using an ECHO fluorescence microscope. Quantification of senescent and apoptotic cells was performed using ImageJ software. Briefly, images were converted to 8-bit, thresholded, binarized, and processed using the watershed function, followed by particle analysis using the ‘Analyze Particles’ function (73). For better visualization of the blue-green precipitate, the images were tinted yellow. All experiments were performed in three independent biological replicates each with technical duplicates.

### RNA isolation

SiHa, SiHa C51S and RPE-1 cells were seeded at a density of 400,000 cells per well in 6-well plates using complete growth medium. The following day, cells were treated with either DMSO (0.02%v/v vehicle control) or 5 µM KTI-240 in 1%FBS media. After 16 hrs, one set of cells was harvested for RNA extraction using the RNeasy Mini Kit (Qiagen, #74104) with on-column DNase digestion (Qiagen, #79254) and processed for RNA sequencing. The second set of cells was collected for protein extraction and Western blotting. Purified RNA samples were submitted for quality control analysis with Agilent’s TapeStation system. All samples had a RIN > 9. The experiment was repeated three independent times.

### RNA Transcriptome analysis

#### Library preparation and sequencing

Indiana University’s Center for Medical Genomics Core processed evaluated the total RNA samples for their quantity and quality using Agilent TapeStation. All the samples had good quality with RIN (RNA Integrity Number) of 9.3-10. One hundred nanograms of total RNA was used for library preparation with the Illumina Stranded mRNA Prep, Ligation kit (lllumina), following the manufacturer’s instruction. Each resulting uniquely dual-indexed library was quantified and quality accessed by Qubit and Agilent TapeStation, and multiple libraries were pooled in equal molarity. The pooled libraries were sequenced with 2×100bp paired-end configuration on an Illumina NovaSeq X PLUS sequencer.

#### RNA-seq data analysis

The sequencing reads were first quality checked using FastQC (v.0.11.5, Babraham Bioinformatics, Cambridge, UK) for quality control. The sequence data were then mapped to the human reference genome hg38 and the Human papillomavirus type 16 (NC_001526.4) using the RNA-seq aligner STAR (v.2.7.10a, (74)) with the following parameter: “--outSAMmapqUnique 60”. To evaluate quality of the RNA-seq data, the number of reads that fell into different annotated regions (exonic, intronic, splicing junction, intergenic, promoter, UTR, etc.) of the reference genome was assessed using bamutils (from ngsutils v.0.4.17, (75)). Uniquely mapped reads were used to quantify the gene level expression employing featureCounts (subread v.2.0.3 (76); with the following parameters: “-s 2 -Q 10”. The data was normalized using TMM (trimmed mean of M values) method. Differential expression analysis was performed in Partek Flow. Features with a maximum ≤ 5.0 were excluded from analysis. Samples were normalized for differential expression analysis with DESeq2 using Median ratio (DESeq2 only) function. Genes with a fold change of ≥ 2, false discovery rate <0.01 and a mean expression ≥ 5 were considered to be significant. Dysregulated signaling pathways were analyzed using Qiagen’s Ingenuity Pathway Analysis platform. In brief, DESeq2 analysis result was uploaded into IPA and processed with the cut-offs FDR<0.01 and a FC of ≥ 2.

### Xenograft studies

Six-week old athymic nude mice were purchased from Envigo (Inotiv). Mice were kept at normal light cycle with free access to water and food. Mice were bred in-house and homozygous male and female nude mice were used in the xenograft studies. Mouse studies were conducted at the LARC animal facility of Indiana University, Indianapolis in accordance with the mouse handling and experimental protocols approved by the Institutional Animal Care and Use Committee (IACUC protocol # 24130). In these studies, 6-9 week old mice were injected subcutaneously in the flank with SiHa cells (2×10^6 cells/mouse), SiHa-luc (1×10^6 cells/mouse), SCC-47 (2×10^6 cells/mouse), SiHa C51S (1×10^6 cells/mouse), or C33a (2×10^6 cells/mouse). Cells were regularly tested and confirmed negative for mycoplasma using Lonza’s MycoAlert® Mycoplasma Detection Assay. Tumors were measured twice a week with calipers and body weight was recorded. Tumor volume was calculated using the formula V = 0.5 * length * width². Unless otherwise stated, drug administration began when tumors reached a size of 50-100 mm^3^. KTI-218 (50 mg/kg) was administered once daily via intraperitoneal (IP) administration in the vehicle consisting of 5% (v/v) NMP, 20% (v/v) Kolliphor RH40, 20% (v/v) PEG400, 55% (v/v) H_2_O. This vehicle was used as vehicle control for all KTI-218 studies. KTI-240 (25 mg/kg) was administered once or twice daily via IP in 5% (v/v) NMP, 20% (v/v) Kolliphor RH40, 75% (v/v) H_2_O, which was used a vehicle control for all KTI-240 experiments. Mice were treated six days a week. Bioluminescence (BLI) of SiHa-luc xenografts was measured at day 8 and day 39 post-cell injection. D-Luciferin (VivoGlo™Luciferin, in vivo grade, P1042) was prepared in 1X phosphate buffered saline (PBS) at a concentration of 200mg/10ml and administered subcutaneously at a single dose of 150 mg/kg. In these studies, animals were anesthetized using isoflurane. Images were acquired over 30 mins with the IVIS SpectrumCT optical imaging system with 15 images taken with a 2 min. delay and a 2 min exposure. After background subtraction, area under the curve (AUC) and Cmax were calculated.

### Pharmacokinetic study

Bioavailability of KTI-218 after a single IP dose of KTI-218 in plasma was investigated in male C57BL/6 mice. KTI-218 was administered to non-fasting mice in 5% NMP + 20% Kolliphor RH 40 + 20% PEG400 + 55% MilliQ water. Mice were euthanized and plasma was collected from three animals at each time point: 0.08, 0.25, 0.5, 1, 2, 4, 8, 12, 16, 24 hours. KTI-218 levels in the murine plasma were analyzed via HPLC-MS/MS.

### Statistical analysis

Densiometric analysis of protein intensities was performed using ImageJ. Target protein expression was normalized to GAPDH and expressed as fold-change. Fluorescence and β-Gal staining intensities were normalized to DMSO control and expressed as fold-change. All normalized data were analyzed with one-way ANOVA with Dunnett’s or Bonferroni post hoc analysis or Students t-test (unpaired) when only two groups were compared. Dose response curves from luciferase and cell viability assays were fitted with a non-linear regression analysis. Tumor growth curves were compared with two-way ANOVA, where a p-value of 0.05 was considered statistically significant. Each experiment was completed at least three independent times (n ≥ 3; * indicates a P< 0.05). All data are expressed as S.E.M unless otherwise stated.

