## Supplementary figures for "Covalent Inhibition of the Human Papillomavirus Type 16 E6 Protein Restores p53 and Suppresses HPV-Driven Tumorigenesis"

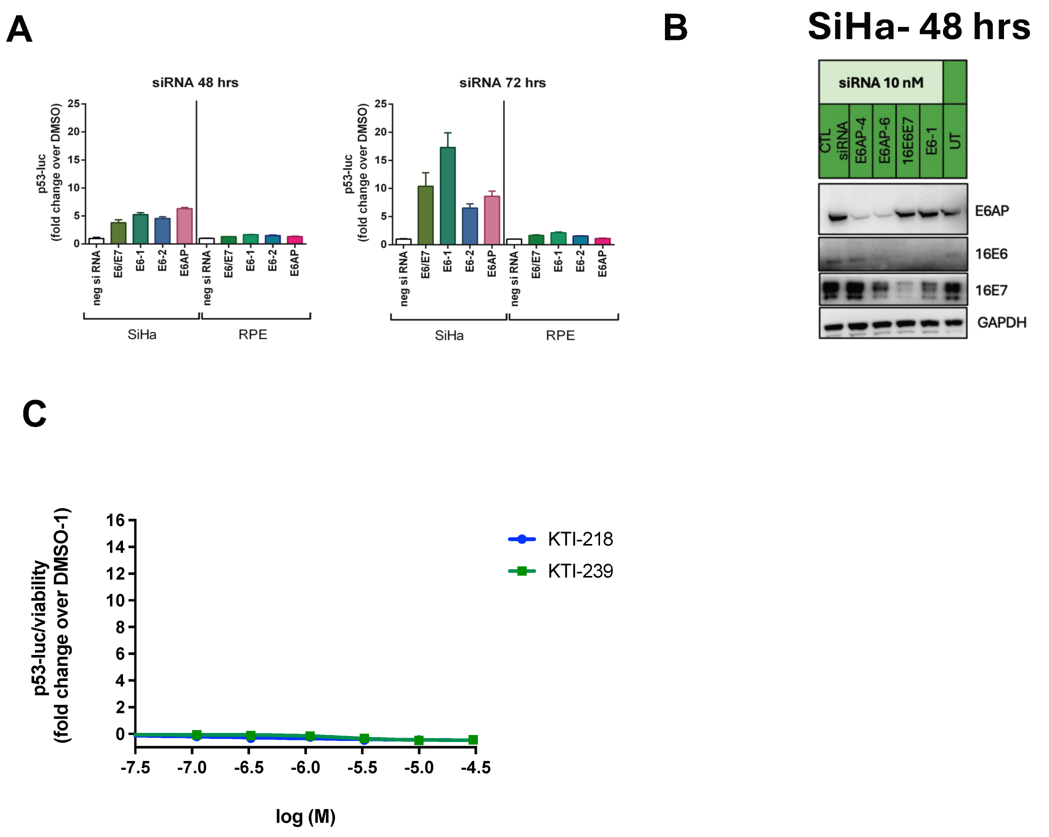


**Supplementary Figure 1**

**(A)** siRNAs (10 nM) targeting 16E6E7, E6 (E6-1, E6-2), E6AP or a scrambled siRNA were transfected into SiHa p53-luc (left) and RPE-1 (right) p53-luc cells. Luc activity was measured 48 or 72 hours after transfection. **(B)** siRNAs were transfected into SiHa. After 48 hours, cells were lysed and E6AP, E6, E7 and GAPDH protein expression were analyzed by Western blot. **(C)** RPE-1 p53-luc cells were exposed to DMSO or increasing concentrations of KTI-218 and KTI-239 for 24 hours.

**A**

| | |
|---|---|
| **Homology Directed Repair (HDR) template** | |
| HPV16_E6_C58S_HDR | GCTGCAAACAACTATACATGATATAATATTAGAATGTGTGT ACTGCAAGCAACAGTTACTGCGACGTGAGGTATATGACTT TGCTTTTCGGGATTTATCAATTGTATACAGAGACGGGAATC CATATGCTGTATGTGATAAATGTTTAAAGTTTTATTCTAAAA TTAGTGAGTA |
| Missense mutations | TGT (C) → TCA (S) |
| Silent mutations | TGTATACAGAGAC |

**B**

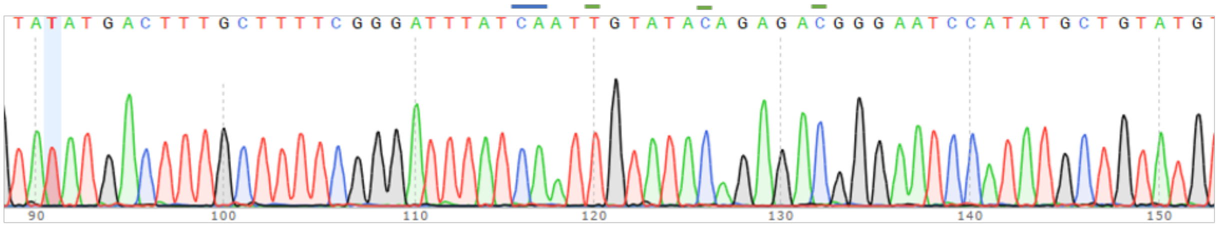


**C**

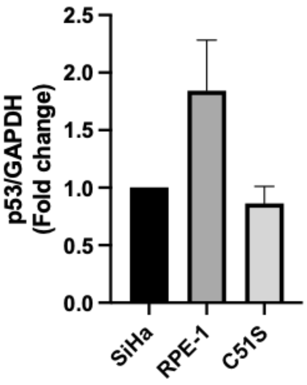


**Supplementary Figure 2**

**(A)** Introduced missense (blue) and silent mutations (green) to generate SiHa 16E6 C51S mutant cells. **(B)** 16E6 sequence with indicated mutations from DNA isolated from SiHa C51S clone. **(C)** Densiometric quantification of p53 protein levels in SiHa, RPE-1 and SiHa C51S cells.

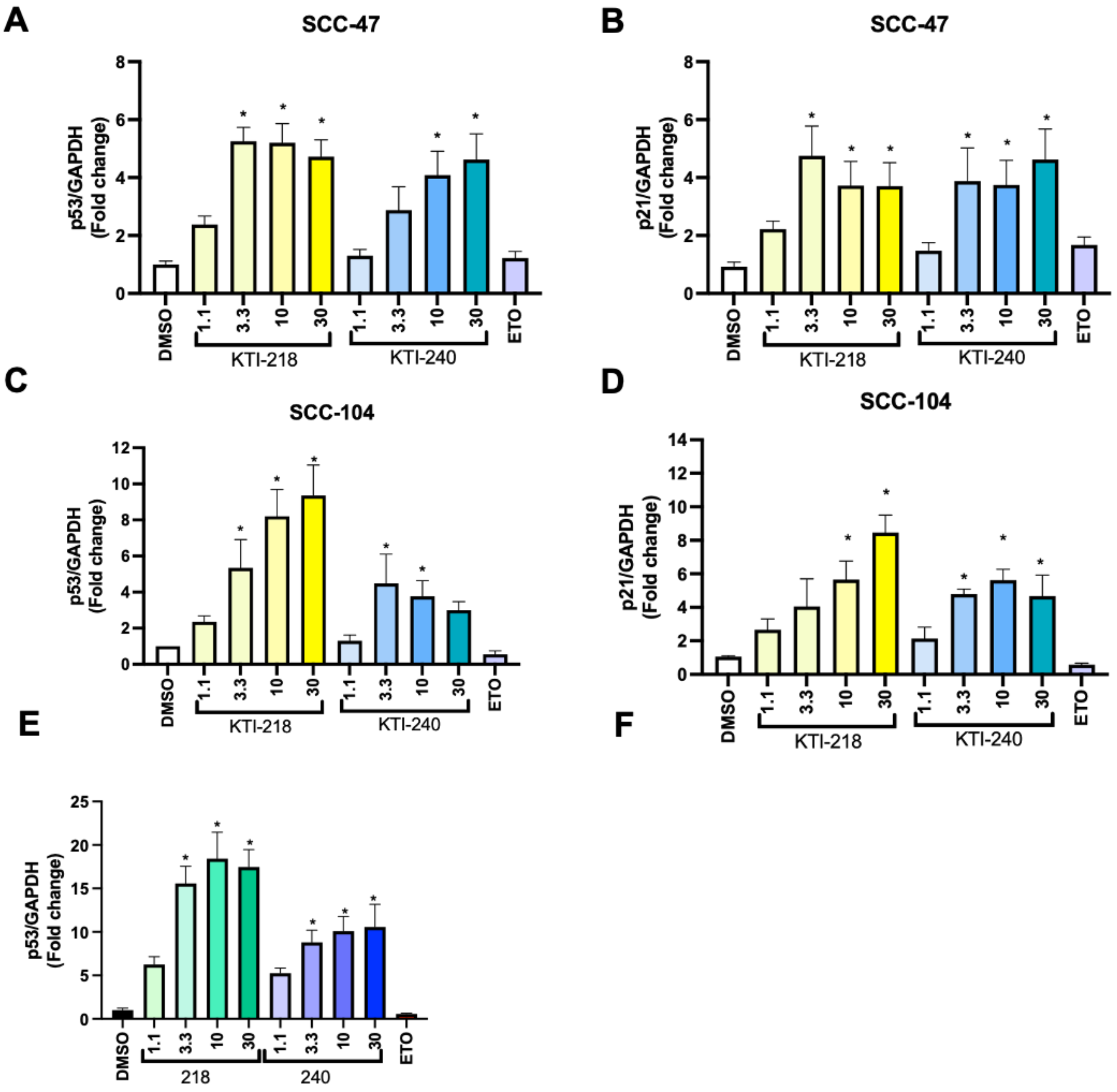


**Supplementary Figure 3**

SCC-47 **(A, B)**, SCC-104 **(C, D)** and W12 cells **(E)** were cultured with KTI-218, KT-240, DMSO (D1, D2) or Etoposide (ETO, 25 µM) for 24 hours, harvested and p53 **(A, C, E)** and p21 **(B, D)** protein levels were analyzed via Immunoblot. GAPDH was used a loading control. Band intensities were quantified by densitometry and normalized to GAPDH / DMSO. Data expressed as S.E.M and each experiment was performed at least three independent times ($n \geq 3$; * indicates $p < 0.05$).

**A**

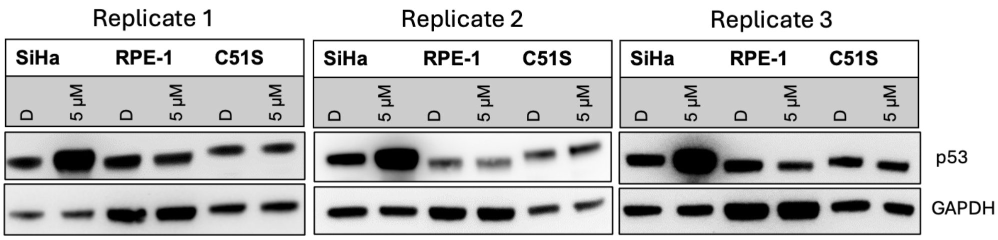


**B**

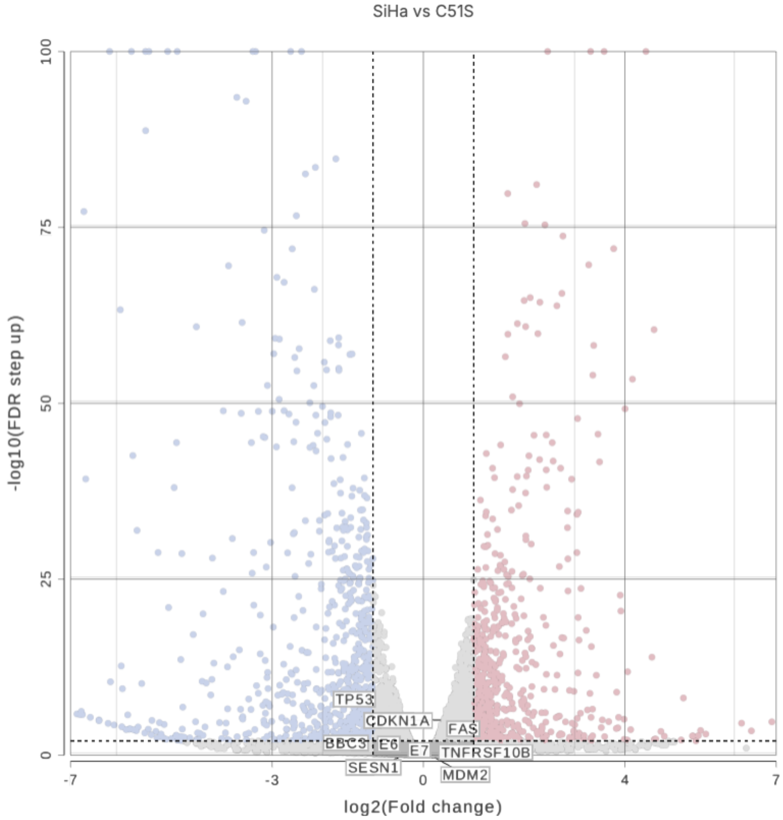


**Supplementary Figure 4**

**(A)** SiHa, SiHa C51S, and RPE-1 cells were cultured in KTI-240 for 16 hours. Lysates from three samples were each immunoblotted to confirm p53 increase. **(B)** RNA was extracted and sequenced from three independent cultures of SiHa and C51S cells to which DMSO at amount equal to the KTI-240 experiment was added. Differentially expressed genes were identified using

DESeq2 analysis with a false discovery rate of <0.01 and a fold change of >2. Volcano plots showing significant transcriptional increases (red) and decreases (blue).

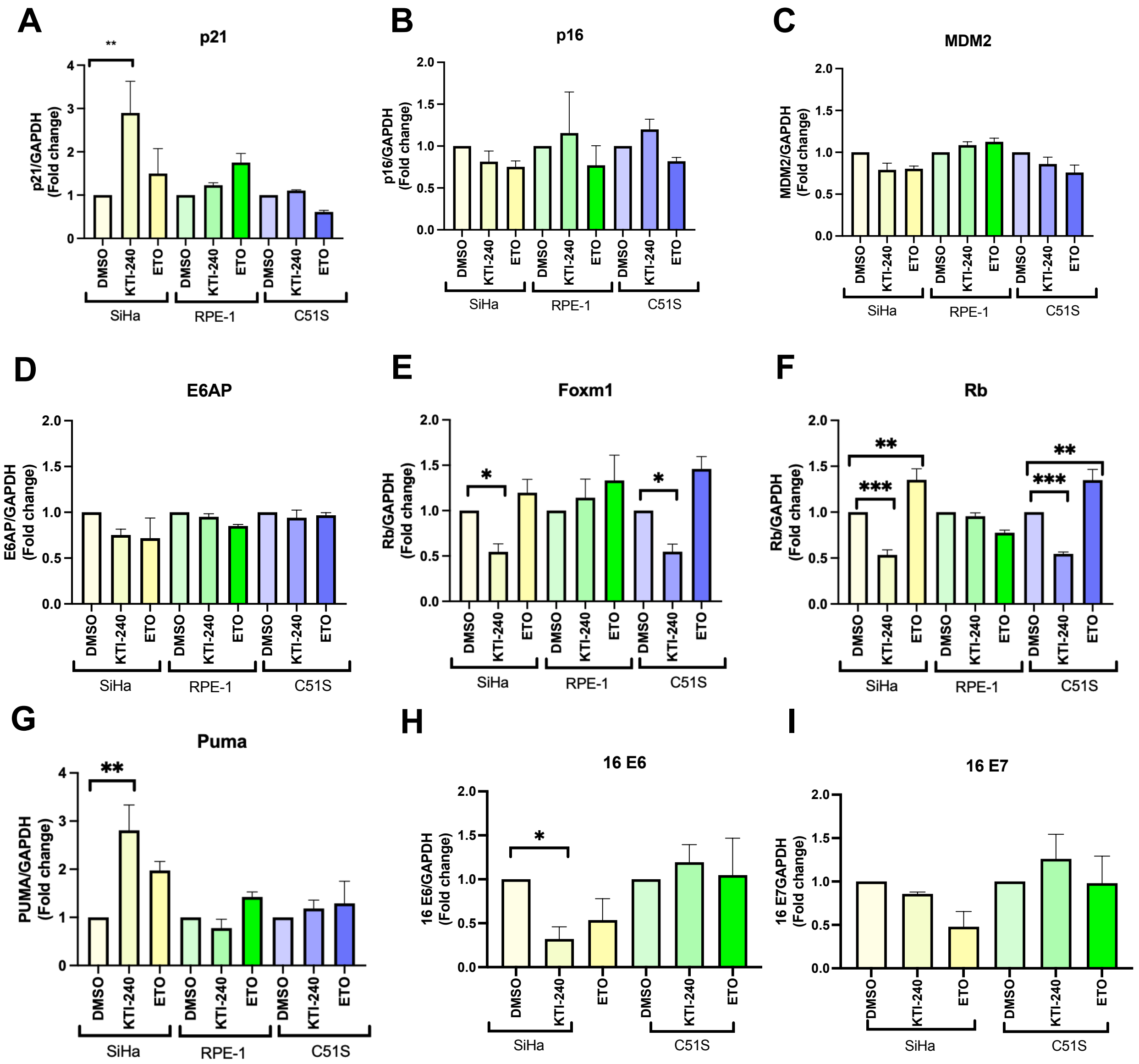


**Supplementary Figure 5**

SiHa, SiHa C51S, and RPE-1 cells were cultured in KTI-240 for 16 hours. Cells were lysed and proteins subjected to Western blot analysis. Band intensities were quantified by densitometry and normalized to GAPDH and compared to DMSO control. Data expressed as S.E.M and each experiment was completed at least three independent times. * indicates a $p < 0.05$).

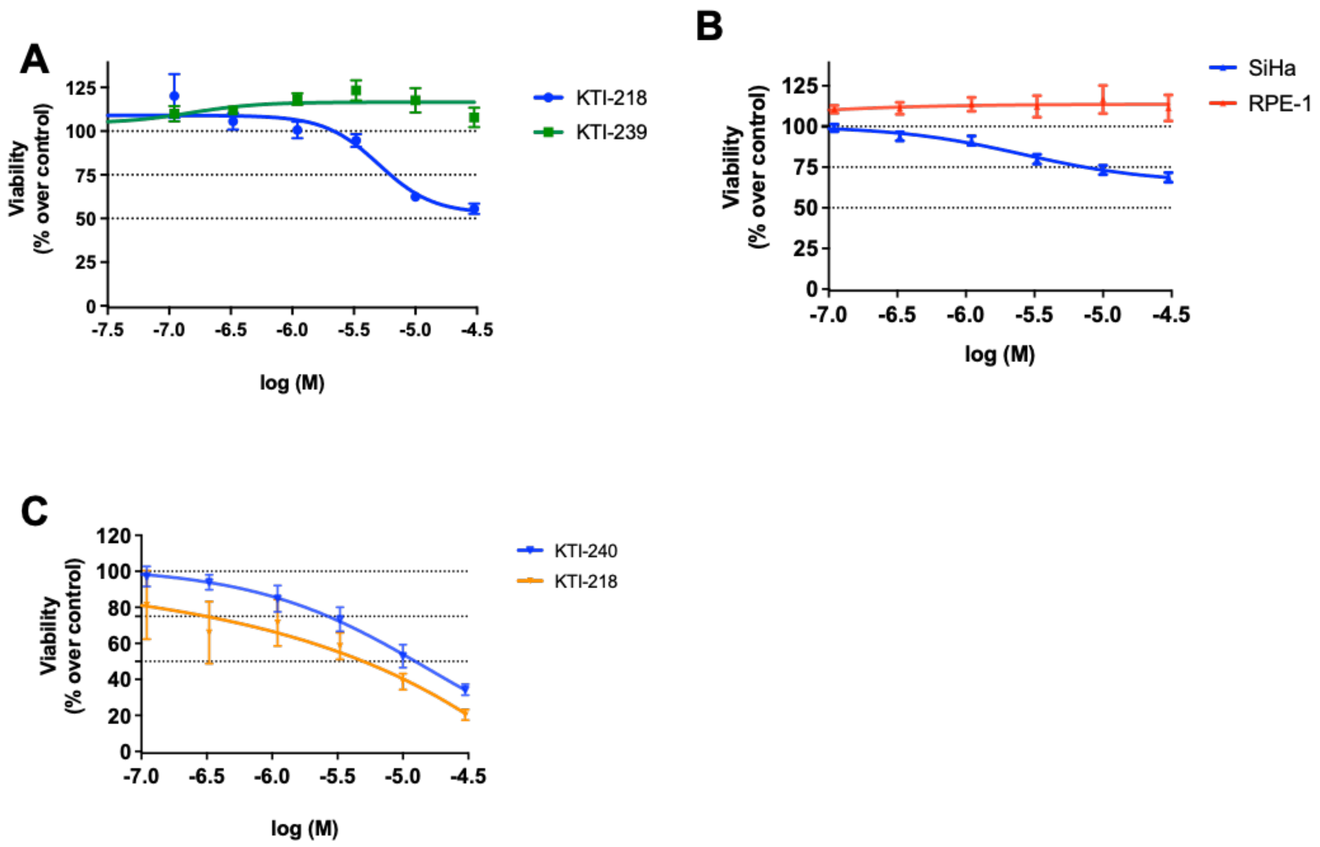


**Supplementary Figure 6**

**(A)** SiHa cells were incubated with increasing concentrations of KTI-218 (blue), non-covalent analog KTI-239 (green) or DMSO for 24 hours. Cell viability was measured using Calcein-AM assay. **(B)** Cell viability of SiHa and RPE-1 cells after 24-hour exposure to KTI-240. **(C)** Viability of W12-E cells was assessed after exposure to increasing doses of KTI-218 or KTI-240 for 24 hours or 48 hours, respectively. Cell viability is expressed as a percent change over DMSO control-treated cells. Data expressed as S.E.M and each experiment was completed at least three independent times.

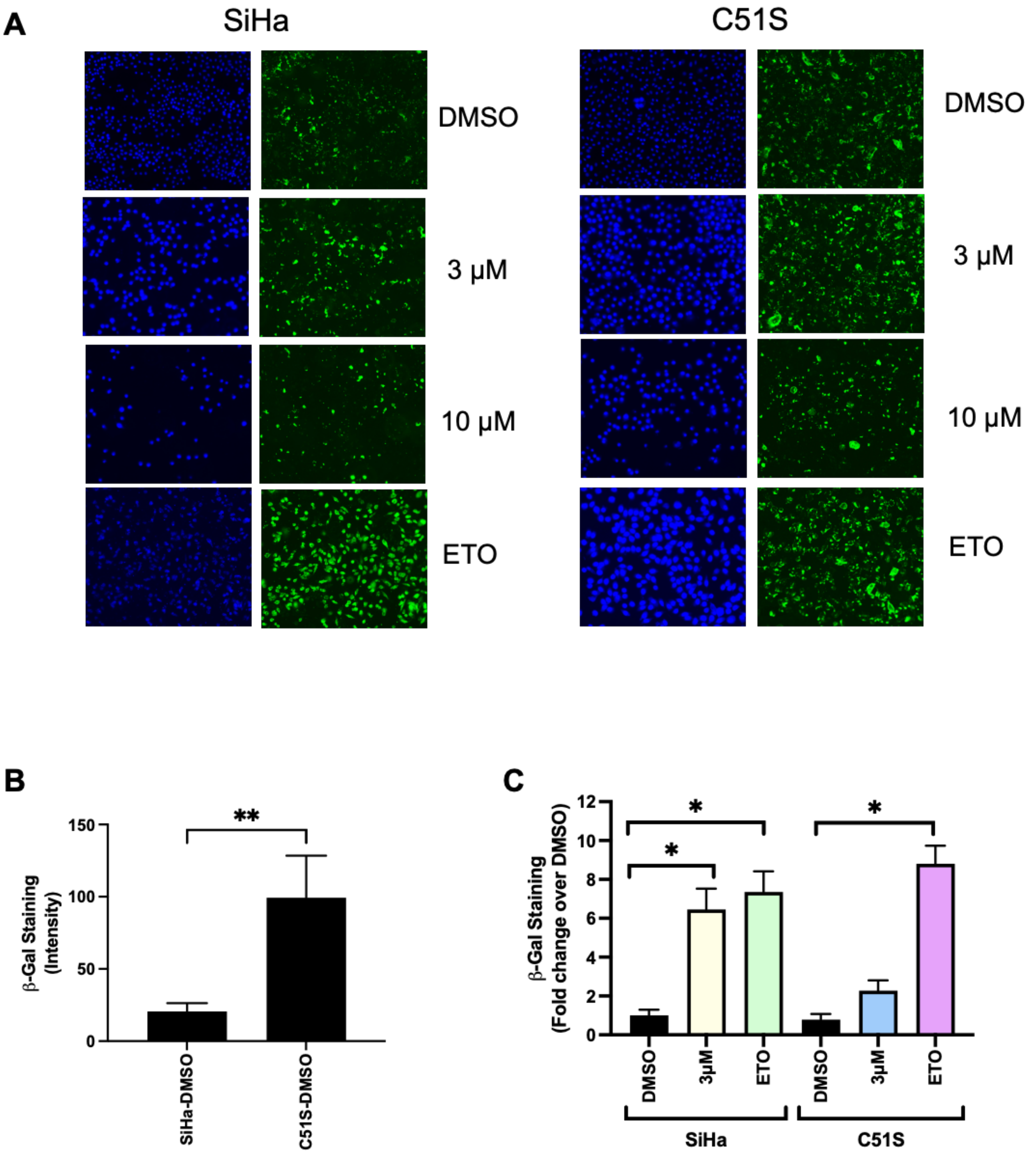


**Supplementary Figure 7**

**(A)** HPV+ cervical SiHa cells (left) and C51S mutant (right) were incubated with DMSO, KTI-240 (3 and 10 μM), or Etoposide (ETO, 2 μM) for 48 hours. Apoptosis was visualized by staining with FITC-Annexin V (Green) and nuclei with DAPI (blue). **(B, C)** Senescence was analyzed via SA-β-gal assay kit after 48 hours of KTI-240 (3 μM), etoposide (ETO, 2 μM) or DMSO treatment. **(B)** Baseline β-gal staining intensities in DMSO treated SiHa and C51S cells. **(C)** β-gal staining intensities after DMSO, KTI-240 and ETO treatment in SiHa and C51S cells. Densiometric analysis was done in ImageJ. Data expressed as S.E.M ($n \geq 3$) and a $p < 0.05$ (*) indicates statistical significance.

**A**

| $AUC_{0\text{-}inf}$ ($h \cdot ng \cdot mL^{-1}$) | $AUC_{0\text{-}tlast}$ ($h \cdot ng \cdot mL^{-1}$) | CL/F ($mL \cdot h^{-1}$) | $C_{last}$ ($ng \cdot mL^{-1}$) | $C_{max}$ ($ng \cdot mL^{-1}$) | $T_{1/2}$ (h) | $k_{el}$ ($h^{-1}$) | $T_{last}$ (h) | $T_{max}$ (h) | $V_d/F$ (mL) |
|---|---|---|---|---|---|---|---|---|---|
| 13786.7 | 13708.2 | 84.86 | 18.39 | 1992.65 | 2.96 | 0.23 | 24 | 1 | 362.28 |

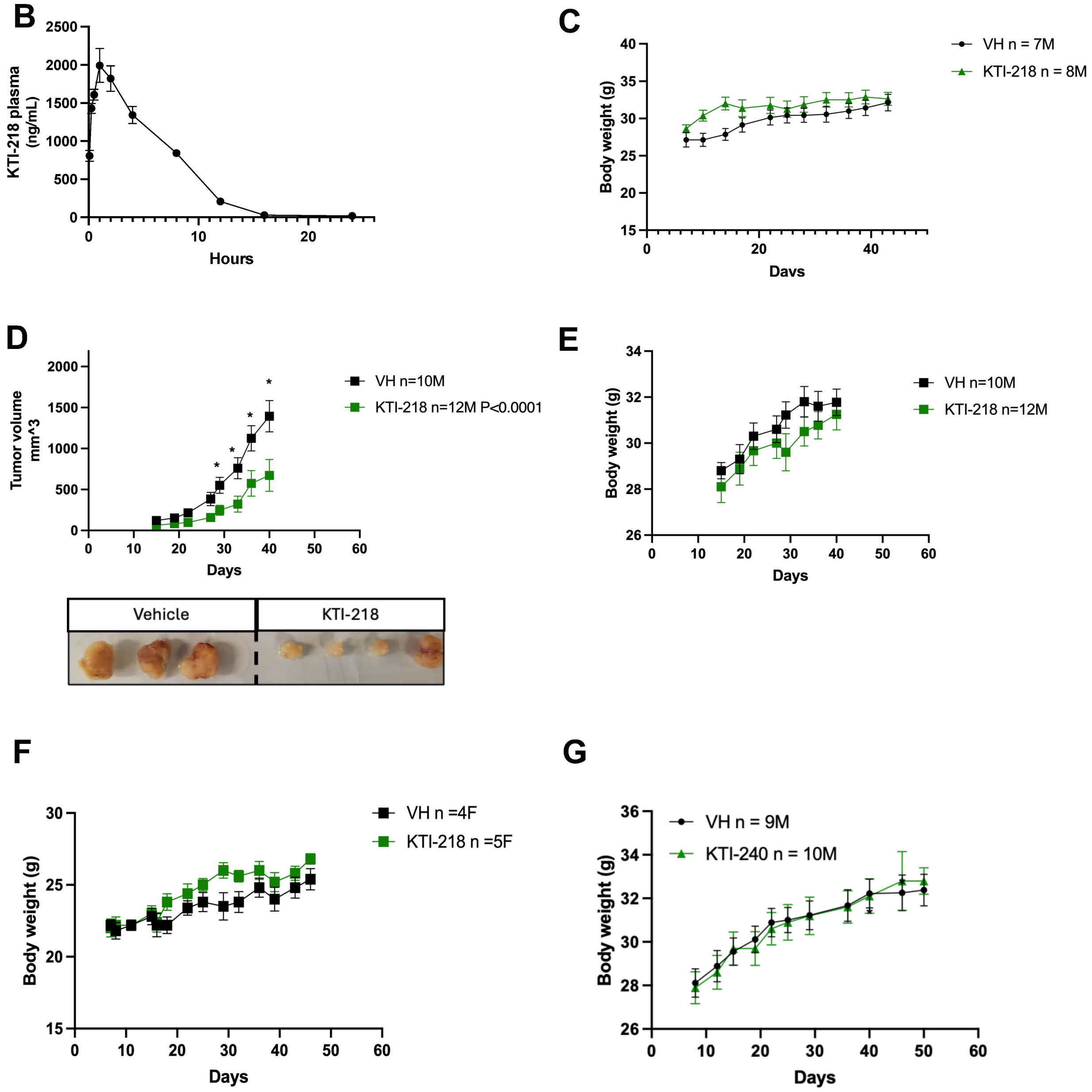

**Supplementary Figure 8**

**(A, B)** KTI-218 plasma levels after a single i.p. injection of 50 mg/kg. **(C)** Body weights of KTI-218 or vehicle treated male nude mice with SCC-47 xenografts. **(D, E)** Male Nu/Nu mice were injected subcutaneously with HPV-16+ SiHa cervical cancer cells. Intraperitoneal injections of 50 mg/kg KTI-218 (n=12, green) or vehicle (VH, n=10 black) began when tumors were ~50-100 $mm^3$. Tumor size was measured by calipers and expressed as S.E.M. and analyzed using two-way ANOVA ($p<0.01$). **(F)** Body weights of KTI-218 or vehicle treated female nude mice with SiHa-luc xenografts. **(G)** Body weights of twice a day intraperitoneally KTI-240 or vehicle treated male nude mice with SCC-47 xenografts.

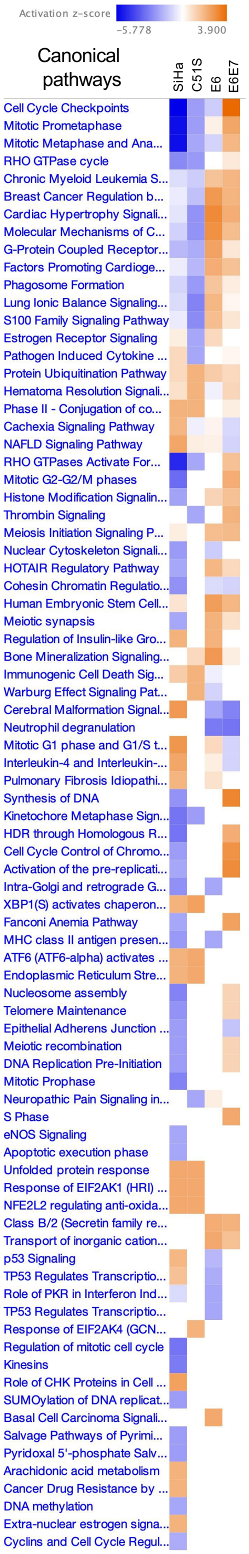
Activation z-score
-5.778
3.900
Canonical pathways
SiHa
C51S
E6
E6E7
Cell Cycle Checkpoints
Mitotic Prometaphase
Mitotic Metaphase and Ana...
RHO GTPase cycle
Chronic Myeloid Leukemia S...
Breast Cancer Regulation b...
Cardiac Hypertrophy Signali...
Molecular Mechanisms of C...
G-Protein Coupled Receptor...
Factors Promoting Cardioge...
Phagosome Formation
Lung Ionic Balance Signaling...
S100 Family Signaling Pathway
Estrogen Receptor Signaling
Pathogen Induced Cytokine ...
Protein Ubiquitination Pathway
Hematoma Resolution Signali...
Phase II - Conjugation of co...
Cachexia Signaling Pathway
NAFLD Signaling Pathway
RHO GTPases Activate For...
Mitotic G2-G2/M phases
Histone Modification Signalin...
Thrombin Signaling
Meiosis Initiation Signaling P...
Nuclear Cytoskeleton Signali...
HOTAIR Regulatory Pathway
Cohesin Chromatin Regulatio...
Human Embryonic Stem Cell...
Meiotic synapsis
Regulation of Insulin-like Gro...
Bone Mineralization Signaling...
Immunogenic Cell Death Sig...
Warburg Effect Signaling Pat...
Cerebral Malformation Signal...
Neutrophil degranulation
Mitotic G1 phase and G1/S t...
Interleukin-4 and Interleukin-...
Pulmonary Fibrosis Idiopathi...
Synthesis of DNA
Kinetochore Metaphase Sign...
HDR through Homologous R...
Cell Cycle Control of Chromo...
Activation of the pre-replicati...
Intra-Golgi and retrograde G...
XBP1(S) activates chaperon...
Fanconi Anemia Pathway
MHC class II antigen presen...
ATF6 (ATF6-alpha) activates ...
Endoplasmic Reticulum Stre...
Nucleosome assembly
Telomere Maintenance
Epithelial Adherens Junction ...
Meiotic recombination
DNA Replication Pre-Initiation
Mitotic Prophase
Neuropathic Pain Signaling in...
S Phase
eNOS Signaling
Apoptotic execution phase
Unfolded protein response
Response of EIF2AK1 (HRI) ...
NFE2L2 regulating anti-oxida...
Class B/2 (Secretin family re...
Transport of inorganic cation...
p53 Signaling
TP53 Regulates Transcriptio...
Role of PKR in Interferon Ind...
TP53 Regulates Transcriptio...
Response of EIF2AK4 (GCN...
Regulation of mitotic cell cycle
Kinesins
Role of CHK Proteins in Cell ...
SUMOylation of DNA replicat...
Basal Cell Carcinoma Signali...
Salvage Pathways of Pyrimi...
Pyridoxal 5'-phosphate Salv...
Arachidonic acid metabolism
Cancer Drug Resistance by ...
DNA methylation
Extra-nuclear estrogen signa...
Cyclins and Cell Cycle Regul...

**Supplementary Figure 9**

IPA Qiagen analysis of differentially affected signaling pathways in KTI-240 cultured SiHa and C51S cells compared to transcriptomes from HNC overexpressing 16E6 or 16E6E7. In the heat map, blue refers to downregulated and orange to upregulated pathways (60).
