## Supplementary Table 1 for "Covalent Inhibition of the Human Papillomavirus Type 16 E6 Protein Restores p53 and Suppresses HPV-Driven Tumorigenesis"

Supplementary Table 1: Differentially expressed genes in SiHa cells after KTI-240 treatment.

| Feature ID | P-value | FDR | Fold change | LSMean(KTI-240) | LSMean(DMSO) |
|---|---|---|---|---|---|
| CDKN1A | 0.00E+00 | 0.00E+00 | 6.94 | 37056.37 | 5341.11 |
| SLC7A11 | 3.23E-202 | 2.60E-198 | 8.91 | 13532.29 | 1519.21 |
| MDM2 | 7.68E-180 | 4.12E-176 | 4.19 | 16898.48 | 4033.53 |
| OSGIN1 | 5.58E-135 | 2.24E-131 | 5.62 | 6938.52 | 1233.52 |
| FDXR | 6.25E-103 | 2.01E-99 | 3.32 | 4647.10 | 1398.54 |
| TXNIP | 2.46E-101 | 6.61E-98 | -3.68 | 1555.82 | 5723.72 |
| RRM2 | 2.83E-97 | 6.52E-94 | -3.19 | 7673.59 | 24457.81 |
| SPATA18 | 1.82E-95 | 3.67E-92 | 5.42 | 1168.22 | 215.63 |
| AURKB | 2.46E-93 | 4.40E-90 | -2.87 | 1288.73 | 3698.97 |
| TK1 | 1.83E-92 | 2.95E-89 | -3.12 | 2352.30 | 7332.17 |
| ZNF469 | 2.82E-91 | 4.13E-88 | 4.87 | 6614.33 | 1358.33 |
| MYBL2 | 1.99E-89 | 2.67E-86 | -2.94 | 4264.12 | 12535.31 |
| MCM7 | 1.59E-83 | 1.97E-80 | -2.40 | 8218.23 | 19750.64 |
| GCLM | 5.08E-83 | 5.84E-80 | 2.39 | 9279.97 | 3881.81 |
| GADD45A | 3.69E-81 | 3.96E-78 | 3.58 | 2384.68 | 665.72 |
| NCAPH | 9.22E-80 | 9.28E-77 | -3.37 | 795.78 | 2679.53 |
| AEN | 3.46E-79 | 3.28E-76 | 2.33 | 10212.52 | 4390.33 |
| SAT1 | 7.39E-79 | 6.61E-76 | 2.74 | 7438.72 | 2714.98 |
| SHCBP1 | 1.66E-78 | 1.41E-75 | -2.41 | 2741.54 | 6605.53 |
| UHRF1 | 3.56E-77 | 2.87E-74 | -2.78 | 1601.51 | 4457.22 |
| PKMYT1 | 1.81E-75 | 1.39E-72 | -2.85 | 1360.75 | 3874.54 |
| BTG2 | 8.65E-74 | 6.33E-71 | 2.99 | 3290.94 | 1101.34 |
| ZWINT | 8.74E-71 | 6.12E-68 | -2.53 | 2423.33 | 6129.71 |
| ASF1B | 3.50E-69 | 2.35E-66 | -2.44 | 2222.90 | 5432.02 |
| PLK1 | 4.39E-68 | 2.82E-65 | -2.42 | 3733.41 | 9030.34 |
| CDC45 | 1.92E-66 | 1.19E-63 | -2.93 | 999.10 | 2926.36 |
| CCNA2 | 3.72E-66 | 2.22E-63 | -2.63 | 1465.52 | 3848.59 |
| GPC1 | 6.54E-66 | 3.76E-63 | 2.25 | 19336.27 | 8606.03 |
| EXO1 | 7.18E-66 | 3.99E-63 | -3.11 | 588.81 | 1831.14 |
| GTSE1 | 8.54E-65 | 4.59E-62 | -2.43 | 1848.92 | 4488.70 |
| E2F2 | 1.39E-64 | 7.20E-62 | -3.64 | 480.07 | 1747.15 |
| KIF18B | 6.41E-64 | 3.22E-61 | -2.67 | 1856.39 | 4957.30 |
| SPC24 | 2.11E-63 | 1.03E-60 | -2.97 | 575.86 | 1710.90 |
| DHRS3 | 1.58E-62 | 7.46E-60 | -2.99 | 802.19 | 2400.73 |
| NMRAL2P | 1.67E-62 | 7.68E-60 | 7.66 | 491.12 | 64.15 |
| CDCA8 | 2.21E-62 | 9.90E-60 | -2.63 | 1960.36 | 5161.55 |
| TP53INP1 | 2.49E-62 | 1.08E-59 | 3.70 | 1860.47 | 503.45 |
| PHLDA3 | 2.72E-62 | 1.15E-59 | 2.55 | 6090.61 | 2392.56 |
| KIF20A | 3.62E-62 | 1.49E-59 | -2.67 | 4165.69 | 11118.41 |
| CDCA3 | 7.55E-62 | 3.04E-59 | -2.69 | 1024.51 | 2756.00 |
| MCM10 | 1.04E-61 | 4.10E-59 | -3.45 | 483.72 | 1669.58 |

Supplementary Table 1: Differentially expressed genes in SiHa cells after KTI-240 treatment.

| | | | | | |
|---|---|---|---|---|---|
| BIRC5 | 2.05E-61 | 7.86E-59 | -2.54 | 2159.67 | 5488.71 |
| LY6D | 1.81E-60 | 6.79E-58 | 3.72 | 3006.82 | 808.32 |
| ACTA2 | 1.92E-60 | 7.03E-58 | 3.10 | 1385.48 | 446.90 |
| NCAPG | 2.17E-60 | 7.78E-58 | -2.59 | 1901.86 | 4920.41 |
| UBE2C | 2.66E-60 | 9.31E-58 | -2.36 | 3731.08 | 8799.36 |
| TNFRSF10B | 2.73E-59 | 9.34E-57 | 2.09 | 11542.62 | 5512.26 |
| BUB1 | 5.97E-59 | 2.00E-56 | -2.36 | 2399.63 | 5667.88 |
| GCLC | 3.23E-58 | 1.06E-55 | 2.70 | 4986.12 | 1849.24 |
| CIT | 5.89E-58 | 1.90E-55 | -2.58 | 2379.39 | 6132.27 |
| MELK | 7.99E-57 | 2.52E-54 | -2.51 | 1261.69 | 3161.68 |
| KIF23 | 4.77E-56 | 1.48E-53 | -2.46 | 4790.53 | 11802.71 |
| FOSL1 | 7.90E-56 | 2.40E-53 | 3.94 | 1809.78 | 458.98 |
| BUB1B | 3.14E-55 | 9.37E-53 | -2.44 | 1783.09 | 4341.99 |
| TRIP13 | 5.45E-55 | 1.60E-52 | -2.14 | 4229.59 | 9033.98 |
| SEC24D | 5.84E-55 | 1.68E-52 | 2.72 | 4495.25 | 1655.48 |
| SQSTM1 | 1.89E-54 | 5.35E-52 | 2.45 | 68957.66 | 28184.39 |
| RAD54L | 2.42E-54 | 6.61E-52 | -2.96 | 548.63 | 1622.92 |
| MXD3 | 1.44E-53 | 3.88E-51 | -2.97 | 384.02 | 1141.64 |
| E2F8 | 1.83E-53 | 4.83E-51 | -3.79 | 241.41 | 915.56 |
| NCAPD2 | 2.19E-53 | 5.69E-51 | -2.14 | 7940.72 | 16954.59 |
| SPAG5 | 5.66E-53 | 1.45E-50 | -2.31 | 4304.65 | 9956.21 |
| RACGAP1 | 2.26E-52 | 5.69E-50 | -2.18 | 3909.83 | 8510.74 |
| MAFG | 9.57E-52 | 2.37E-49 | 2.29 | 6388.66 | 2791.91 |
| SESN1 | 2.51E-51 | 6.13E-49 | 2.36 | 2152.63 | 911.51 |
| TACC3 | 5.07E-51 | 1.22E-48 | -2.35 | 2295.78 | 5400.79 |
| SULF2 | 5.78E-51 | 1.37E-48 | 8.03 | 402.12 | 50.09 |
| RFC3 | 6.03E-51 | 1.41E-48 | -2.29 | 1200.49 | 2749.11 |
| FAS | 1.94E-50 | 4.47E-48 | 2.52 | 1888.17 | 749.25 |
| EPGN | 2.56E-50 | 5.81E-48 | 4.91 | 854.95 | 174.22 |
| FOXM1 | 5.61E-50 | 1.25E-47 | -2.14 | 4022.36 | 8615.49 |
| TNFAIP2 | 1.11E-49 | 2.46E-47 | -2.48 | 6613.93 | 16429.23 |
| KRT80 | 1.78E-49 | 3.86E-47 | -2.93 | 360.63 | 1057.85 |
| F3 | 5.27E-49 | 1.13E-46 | 2.63 | 4894.33 | 1858.44 |
| CDC20 | 6.37E-49 | 1.35E-46 | -2.22 | 6083.77 | 13495.80 |
| PRC1 | 1.01E-48 | 2.10E-46 | -2.30 | 7445.93 | 17109.74 |
| MASTL | 2.24E-48 | 4.62E-46 | -2.41 | 973.16 | 2346.63 |
| CDCA5 | 2.85E-48 | 5.80E-46 | -2.44 | 1530.71 | 3727.56 |
| DNAJB9 | 3.34E-48 | 6.72E-46 | 3.05 | 1686.82 | 552.67 |
| XBP1 | 5.67E-48 | 1.13E-45 | 2.11 | 5565.39 | 2636.37 |
| TOP2A | 3.05E-47 | 5.99E-45 | -2.35 | 8510.09 | 19977.95 |
| SESN2 | 4.84E-46 | 9.17E-44 | 2.28 | 2785.52 | 1219.36 |
| SRXN1 | 7.75E-46 | 1.43E-43 | 2.55 | 13024.82 | 5102.05 |
| CCNB2 | 1.96E-45 | 3.58E-43 | -2.22 | 2895.45 | 6440.29 |

Supplementary Table 1:  Differentially expressed genes in SiHa cells after KTI-240 treatment.

| | | | | | |
|---|---|---|---|---|---|
| H1F0 | 2.03E-45 | 3.67E-43 | -2.14 | 10291.75 | 21999.33 |
| HSP90B1 | 5.65E-45 | 1.01E-42 | 2.23 | 47155.02 | 21121.57 |
| TNFSF10 | 3.66E-44 | 6.47E-42 | -3.04 | 318.41 | 967.84 |
| TNS3 | 4.03E-44 | 7.05E-42 | -2.05 | 6353.98 | 13006.53 |
| TTK | 7.60E-44 | 1.32E-41 | -2.49 | 605.79 | 1511.24 |
| HJURP | 1.83E-43 | 3.13E-41 | -2.21 | 1185.15 | 2616.85 |
| ERCC6L | 1.94E-43 | 3.29E-41 | -2.81 | 385.92 | 1084.10 |
| PAPPA | 5.49E-43 | 9.17E-41 | 2.69 | 1480.70 | 550.19 |
| KIF15 | 5.53E-43 | 9.17E-41 | -2.56 | 821.74 | 2102.98 |
| PBK | 7.00E-43 | 1.15E-40 | -2.43 | 791.95 | 1926.23 |
| ANLN | 1.37E-42 | 2.23E-40 | -2.66 | 4278.94 | 11367.61 |
| C11orf86 | 1.42E-42 | 2.29E-40 | -2.34 | 1276.18 | 2989.92 |
| EREG | 2.19E-42 | 3.49E-40 | 2.07 | 18705.86 | 9025.59 |
| ATF3 | 2.84E-42 | 4.49E-40 | 3.55 | 943.35 | 265.79 |
| KNL1 | 4.21E-42 | 6.58E-40 | -2.24 | 1434.24 | 3211.59 |
| UGDH | 5.06E-42 | 7.83E-40 | 2.47 | 10276.51 | 4161.86 |
| DEPDC1 | 6.67E-42 | 1.02E-39 | -2.22 | 1927.11 | 4285.90 |
| CENPI | 7.86E-42 | 1.18E-39 | -2.21 | 765.22 | 1693.71 |
| ESPL1 | 9.09E-42 | 1.36E-39 | -2.06 | 1476.29 | 3038.90 |
| PSRC1 | 1.04E-41 | 1.53E-39 | -2.35 | 822.13 | 1931.58 |
| VSIR | 1.90E-41 | 2.78E-39 | -2.57 | 686.07 | 1761.40 |
| KIF2C | 2.43E-41 | 3.52E-39 | -2.22 | 3414.50 | 7575.82 |
| NDC80 | 1.01E-40 | 1.46E-38 | -2.82 | 743.14 | 2099.36 |
| ALDH3A1 | 1.51E-40 | 2.13E-38 | 2.57 | 8202.12 | 3185.82 |
| HSPA5 | 1.65E-40 | 2.32E-38 | 2.55 | 68540.44 | 26911.27 |
| PLK3 | 6.00E-40 | 8.33E-38 | 2.54 | 1351.74 | 531.33 |
| PLK4 | 7.63E-40 | 1.05E-37 | -2.45 | 522.22 | 1280.69 |
| UBE2T | 9.65E-40 | 1.32E-37 | -2.10 | 1228.41 | 2577.91 |
| SYNPO | 3.91E-39 | 5.29E-37 | -2.38 | 1666.34 | 3959.57 |
| DEPDC1B | 5.89E-39 | 7.83E-37 | -2.37 | 782.78 | 1855.73 |
| CKB | 8.99E-39 | 1.19E-36 | -2.12 | 2721.51 | 5776.44 |
| AKR1C2 | 1.79E-38 | 2.35E-36 | 2.54 | 51683.14 | 20362.33 |
| KIF4A | 2.21E-38 | 2.86E-36 | -2.28 | 2312.25 | 5276.82 |
| LMNB1 | 2.29E-38 | 2.93E-36 | -2.01 | 4331.20 | 8723.82 |
| INCENP | 2.29E-38 | 2.93E-36 | -2.16 | 2544.48 | 5490.29 |
| FANCD2 | 4.95E-38 | 6.28E-36 | -2.05 | 1549.94 | 3173.37 |
| RAD51 | 1.19E-37 | 1.49E-35 | -2.39 | 526.32 | 1255.80 |
| ABCC1 | 2.43E-37 | 3.03E-35 | 2.23 | 1530.48 | 687.42 |
| RAD51AP1 | 4.78E-37 | 5.87E-35 | -2.56 | 556.01 | 1421.17 |
| TPX2 | 8.91E-37 | 1.08E-34 | -2.38 | 7167.95 | 17078.24 |
| CPA4 | 1.16E-36 | 1.40E-34 | -3.75 | 132.73 | 497.79 |
| ORC6 | 2.12E-36 | 2.53E-34 | -2.03 | 1325.40 | 2693.54 |
| NUSAP1 | 7.26E-36 | 8.60E-34 | -2.38 | 3823.85 | 9086.20 |

Supplementary Table 1: Differentially expressed genes in SiHa cells after KTI-240 treatment.

| | | | | | |
|---|---|---|---|---|---|
| CENPM | 8.15E-36 | 9.58E-34 | -2.10 | 835.27 | 1758.14 |
| SKA3 | 9.79E-36 | 1.13E-33 | -2.73 | 292.97 | 798.95 |
| CCNF | 1.29E-35 | 1.49E-33 | -2.07 | 2522.40 | 5230.53 |
| GPRC5A | 1.51E-35 | 1.72E-33 | -2.20 | 5227.62 | 11501.14 |
| MAD2L1 | 1.73E-35 | 1.96E-33 | -2.20 | 967.70 | 2131.93 |
| KLF4 | 2.06E-35 | 2.32E-33 | 2.63 | 927.45 | 353.31 |
| DTL | 2.62E-35 | 2.93E-33 | -2.02 | 1483.94 | 2997.73 |
| ECT2 | 2.69E-35 | 2.98E-33 | -2.08 | 3434.77 | 7136.24 |
| TNFRSF10D | 3.03E-35 | 3.34E-33 | 2.25 | 1407.00 | 626.06 |
| CEP55 | 1.54E-34 | 1.68E-32 | -2.13 | 1629.14 | 3474.12 |
| PANX2 | 1.78E-34 | 1.92E-32 | 2.44 | 1098.93 | 450.18 |
| NEK2 | 1.95E-34 | 2.10E-32 | -2.28 | 911.84 | 2077.11 |
| PRR11 | 3.18E-34 | 3.39E-32 | -2.08 | 2331.67 | 4859.45 |
| MKI67 | 6.70E-34 | 6.96E-32 | -2.37 | 8249.28 | 19563.30 |
| CDCA2 | 9.99E-34 | 1.03E-31 | -2.23 | 1025.70 | 2290.73 |
| SLC12A3 | 1.01E-33 | 1.04E-31 | -2.05 | 8588.04 | 17626.49 |
| BBC3 | 1.23E-33 | 1.24E-31 | 2.72 | 1320.92 | 486.04 |
| SAPCD2 | 2.00E-33 | 2.00E-31 | -2.05 | 2647.57 | 5437.66 |
| GINS4 | 2.79E-33 | 2.76E-31 | -2.05 | 847.99 | 1740.34 |
| BLM | 3.24E-33 | 3.19E-31 | -2.38 | 527.66 | 1254.91 |
| DLGAP5 | 3.34E-33 | 3.26E-31 | -2.47 | 1352.28 | 3337.30 |
| BICDL1 | 5.22E-33 | 5.06E-31 | 2.26 | 2373.27 | 1051.96 |
| CDC25C | 6.18E-33 | 5.96E-31 | -2.39 | 772.28 | 1846.27 |
| FAM111B | 1.22E-32 | 1.16E-30 | -2.65 | 728.98 | 1931.60 |
| GAS2L3 | 2.03E-32 | 1.91E-30 | -2.21 | 527.49 | 1164.40 |
| NUCB2 | 2.23E-32 | 2.08E-30 | 2.01 | 4015.67 | 1997.06 |
| HMGB2 | 2.28E-32 | 2.13E-30 | -2.20 | 3505.58 | 7728.74 |
| CDK1 | 3.32E-32 | 3.07E-30 | -2.46 | 1984.00 | 4871.12 |
| ESCO2 | 4.89E-32 | 4.50E-30 | -3.02 | 349.54 | 1056.93 |
| FANCI | 1.31E-31 | 1.19E-29 | -2.08 | 3900.33 | 8113.21 |
| PIMREG | 2.58E-31 | 2.30E-29 | -2.21 | 922.54 | 2034.59 |
| MIR34AHG | 3.86E-31 | 3.44E-29 | 2.37 | 985.29 | 415.94 |
| MANF | 7.78E-31 | 6.88E-29 | 2.02 | 6185.66 | 3064.88 |
| CDKN3 | 8.59E-31 | 7.56E-29 | -2.13 | 1003.29 | 2135.62 |
| IL7R | 9.16E-31 | 8.02E-29 | -2.87 | 299.52 | 859.71 |
| LIF | 1.26E-30 | 1.09E-28 | 3.04 | 680.48 | 223.89 |
| SERPINE1 | 1.30E-30 | 1.12E-28 | 3.38 | 635.32 | 188.24 |
| TNFRSF21 | 1.47E-30 | 1.26E-28 | 2.55 | 865.48 | 339.91 |
| WIPI1 | 2.24E-30 | 1.90E-28 | 2.15 | 1465.57 | 680.72 |
| CENPA | 3.01E-30 | 2.53E-28 | -2.15 | 647.64 | 1391.71 |
| TJP3 | 3.68E-30 | 3.05E-28 | -2.38 | 336.02 | 798.59 |
| SLC6A9 | 4.61E-30 | 3.76E-28 | 2.01 | 2533.00 | 1257.37 |
| SGO1 | 6.62E-30 | 5.33E-28 | -2.39 | 378.83 | 904.95 |

Supplementary Table 1: Differentially expressed genes in SiHa cells after KTI-240 treatment.

| | | | | | |
|---|---|---|---|---|---|
| GPAT3 | 1.33E-29 | 1.07E-27 | 2.66 | 555.81 | 209.25 |
| SNAI1 | 1.61E-29 | 1.27E-27 | 2.00 | 2360.05 | 1178.94 |
| FBXO5 | 4.98E-29 | 3.89E-27 | -2.34 | 576.31 | 1348.23 |
| IGSF10 | 5.51E-29 | 4.29E-27 | -2.29 | 541.57 | 1238.63 |
| PLPP5 | 9.45E-29 | 7.24E-27 | 2.23 | 2196.45 | 983.63 |
| KIF11 | 1.40E-28 | 1.06E-26 | -2.33 | 2197.24 | 5123.94 |
| HYOU1 | 1.58E-28 | 1.18E-26 | 2.00 | 22656.58 | 11306.21 |
| TICRR | 1.89E-28 | 1.40E-26 | -2.11 | 2352.59 | 4973.42 |
| HMOX1 | 1.29E-27 | 9.32E-26 | 8.58 | 65570.68 | 7643.10 |
| PIF1 | 2.56E-27 | 1.83E-25 | -2.17 | 512.10 | 1111.90 |
| WISP2 | 2.79E-27 | 1.97E-25 | -4.93 | 43.72 | 215.72 |
| FAM114A1 | 3.05E-27 | 2.15E-25 | 2.01 | 1113.02 | 553.50 |
| ZNF367 | 4.41E-27 | 3.09E-25 | -2.16 | 747.87 | 1612.41 |
| CBLB | 6.69E-27 | 4.64E-25 | 2.26 | 1421.70 | 627.75 |
| MTFR2 | 2.96E-26 | 1.98E-24 | -2.89 | 158.26 | 458.07 |
| AMIGO2 | 3.84E-26 | 2.54E-24 | 2.23 | 2239.55 | 1002.56 |
| NRP2 | 4.25E-26 | 2.80E-24 | -3.58 | 107.59 | 384.75 |
| SLC48A1 | 1.06E-25 | 6.92E-24 | 2.12 | 3115.74 | 1471.78 |
| TEDC1 | 2.46E-25 | 1.57E-23 | -2.18 | 474.17 | 1033.85 |
| CLCA2 | 8.64E-25 | 5.41E-23 | 7.88 | 154.25 | 19.58 |
| AKR1C1 | 1.61E-24 | 9.96E-23 | 2.16 | 13856.77 | 6419.71 |
| CKAP2L | 1.65E-24 | 1.02E-22 | -2.31 | 729.41 | 1688.15 |
| CLSPN | 8.42E-24 | 5.06E-22 | -2.11 | 1024.73 | 2157.37 |
| NREP | 1.10E-23 | 6.56E-22 | -2.26 | 413.11 | 934.36 |
| HASPIN | 1.23E-23 | 7.28E-22 | -2.08 | 375.34 | 780.67 |
| DIAPH3 | 1.78E-23 | 1.04E-21 | -2.27 | 664.71 | 1507.83 |
| MAP2K6 | 2.16E-23 | 1.25E-21 | -2.28 | 406.38 | 927.03 |
| CENPU | 2.27E-23 | 1.31E-21 | -2.29 | 351.38 | 805.73 |
| AOX1 | 5.35E-23 | 3.03E-21 | -2.23 | 3245.07 | 7235.66 |
| TSKU | 1.12E-22 | 6.33E-21 | 2.01 | 3745.95 | 1866.36 |
| KIF24 | 2.35E-22 | 1.30E-20 | -2.11 | 456.26 | 960.73 |
| DUSP4 | 3.51E-22 | 1.93E-20 | 2.28 | 706.65 | 310.24 |
| DBP | 3.83E-22 | 2.10E-20 | -2.28 | 307.14 | 700.86 |
| CHAC1 | 1.03E-21 | 5.56E-20 | 4.11 | 451.36 | 109.84 |
| SPRY4 | 1.45E-21 | 7.80E-20 | 4.34 | 404.03 | 93.10 |
| HIRIP3 | 3.37E-21 | 1.77E-19 | -2.14 | 492.42 | 1052.49 |
| ISYNA1 | 4.28E-21 | 2.25E-19 | 2.02 | 2576.85 | 1278.69 |
| TRIM22 | 4.45E-21 | 2.32E-19 | 2.54 | 591.83 | 233.24 |
| SLC37A3 | 5.36E-20 | 2.65E-18 | 2.04 | 963.33 | 471.64 |
| PKN3 | 7.43E-20 | 3.63E-18 | -2.13 | 520.86 | 1106.82 |
| CABYR | 1.58E-19 | 7.62E-18 | 2.02 | 3105.29 | 1537.33 |
| CPEB2 | 3.48E-19 | 1.66E-17 | 2.42 | 718.43 | 296.57 |
| AQP3 | 8.22E-19 | 3.85E-17 | -2.05 | 1998.13 | 4086.92 |

Supplementary Table 1: Differentially expressed genes in SiHa cells after KTI-240 treatment.

| | | | | | |
|---|---|---|---|---|---|
| MMP3 | 1.24E-18 | 5.75E-17 | 10.17 | 101.98 | 10.02 |
| SLC29A1 | 1.43E-18 | 6.59E-17 | -2.02 | 368.90 | 743.69 |
| ACER2 | 2.29E-18 | 1.05E-16 | 4.52 | 161.72 | 35.79 |
| NRCAM | 1.33E-17 | 5.81E-16 | 3.97 | 162.51 | 40.95 |
| GDF15 | 2.47E-17 | 1.05E-15 | 30.13 | 699.81 | 23.23 |
| LURAP1L | 2.70E-17 | 1.14E-15 | 2.77 | 485.49 | 175.18 |
| ABCC2 | 3.24E-17 | 1.36E-15 | 2.32 | 432.46 | 186.74 |
| SH3PXD2A | 3.34E-17 | 1.40E-15 | -2.35 | 216.79 | 509.77 |
| CYP4F11 | 1.03E-16 | 4.12E-15 | 9.82 | 121.29 | 12.35 |
| TMPO-AS1 | 1.18E-16 | 4.72E-15 | -2.26 | 342.25 | 772.72 |
| AFF3 | 1.26E-16 | 5.00E-15 | -2.69 | 143.21 | 385.20 |
| ARHGEF39 | 1.60E-16 | 6.35E-15 | -2.04 | 237.52 | 485.22 |
| SLC22A4 | 2.15E-16 | 8.43E-15 | 2.05 | 597.59 | 291.07 |
| MIR1204 | 4.56E-16 | 1.74E-14 | 4.89 | 140.47 | 28.70 |
| METTL7A | 5.92E-16 | 2.23E-14 | -2.58 | 131.65 | 339.39 |
| FICD | 6.82E-16 | 2.55E-14 | 2.09 | 565.52 | 271.22 |
| RTKN2 | 2.14E-15 | 7.76E-14 | -2.45 | 142.04 | 347.50 |
| KIAA1324 | 2.92E-15 | 1.04E-13 | 3.67 | 209.43 | 57.13 |
| NUF2 | 4.90E-15 | 1.72E-13 | -2.01 | 472.41 | 948.97 |
| OLFML2B | 7.33E-15 | 2.55E-13 | -4.10 | 36.95 | 151.66 |
| MAP1B | 1.22E-14 | 4.20E-13 | 2.05 | 602.09 | 293.30 |
| F2R | 1.62E-14 | 5.48E-13 | 2.22 | 363.68 | 163.50 |
| COL17A1 | 2.40E-14 | 7.98E-13 | 4.13 | 128.70 | 31.19 |
| HIST1H2BJ | 2.49E-14 | 8.26E-13 | -2.09 | 506.56 | 1056.73 |
| TUBB4A | 3.05E-14 | 1.00E-12 | -3.45 | 48.49 | 167.49 |
| TVP23C | 4.68E-14 | 1.51E-12 | 2.51 | 492.11 | 196.23 |
| LOC1053712 | 7.30E-14 | 2.31E-12 | 3.18 | 180.14 | 56.64 |
| BTBD19 | 9.06E-14 | 2.85E-12 | 3.40 | 160.56 | 47.27 |
| NOG | 1.16E-13 | 3.62E-12 | -2.18 | 155.77 | 339.53 |
| GPRC5C | 1.30E-13 | 3.99E-12 | 2.54 | 287.17 | 112.93 |
| ENC1 | 1.31E-13 | 4.04E-12 | 2.15 | 520.23 | 241.42 |
| CYP4F3 | 1.67E-13 | 5.11E-12 | 3.63 | 242.25 | 66.81 |
| GJA5 | 1.70E-13 | 5.18E-12 | -3.45 | 57.82 | 199.20 |
| VEPH1 | 1.86E-13 | 5.60E-12 | -2.78 | 89.82 | 249.83 |
| HMMR | 2.11E-13 | 6.35E-12 | -2.07 | 652.03 | 1352.39 |
| ELF3 | 3.38E-13 | 9.86E-12 | -5.78 | 15.82 | 91.46 |
| EPHA4 | 1.36E-12 | 3.75E-11 | -3.07 | 61.52 | 188.78 |
| EDN2 | 1.84E-12 | 5.01E-11 | 3.35 | 149.30 | 44.50 |
| PURPL | 1.88E-12 | 5.12E-11 | 2.83 | 184.55 | 65.21 |
| EME1 | 2.23E-12 | 5.97E-11 | -2.09 | 266.63 | 558.17 |
| HES2 | 2.54E-12 | 6.77E-11 | 2.90 | 157.33 | 54.27 |
| TMEM52B | 2.98E-12 | 7.87E-11 | -2.44 | 94.68 | 230.64 |
| MEIOB | 4.73E-12 | 1.22E-10 | -2.06 | 156.29 | 322.34 |

Supplementary Table 1: Differentially expressed genes in SiHa cells after KTI-240 treatment.

| | | | | | |
|---|---|---|---|---|---|
| NEIL3 | 5.13E-12 | 1.32E-10 | -2.04 | 263.54 | 538.27 |
| KCNS1 | 5.24E-12 | 1.34E-10 | -4.77 | 19.04 | 90.79 |
| GABRQ | 1.69E-11 | 4.10E-10 | -2.24 | 109.90 | 246.54 |
| SCN3B | 3.38E-11 | 8.06E-10 | 2.92 | 148.96 | 50.95 |
| SPC25 | 7.97E-11 | 1.83E-09 | -4.17 | 25.71 | 107.27 |
| ABCA9 | 9.42E-11 | 2.15E-09 | 2.16 | 519.04 | 240.47 |
| PRLR | 1.13E-10 | 2.54E-09 | -3.82 | 26.11 | 99.67 |
| KRT4 | 1.30E-10 | 2.88E-09 | -2.01 | 202.44 | 406.13 |
| ADM2 | 1.48E-10 | 3.27E-09 | 7.12 | 442.48 | 62.18 |
| LINC01260 | 1.99E-10 | 4.35E-09 | -4.70 | 18.85 | 88.64 |
| GRB10 | 2.03E-10 | 4.42E-09 | 2.24 | 296.84 | 132.60 |
| NCCRP1 | 3.32E-10 | 7.13E-09 | -2.60 | 63.89 | 166.29 |
| C3orf80 | 3.63E-10 | 7.75E-09 | 2.03 | 369.86 | 182.43 |
| PGBD5 | 4.29E-10 | 9.07E-09 | 2.37 | 197.18 | 83.24 |
| ABCA17P | 6.20E-10 | 1.28E-08 | -2.12 | 130.17 | 276.25 |
| MAMDC4 | 6.69E-10 | 1.37E-08 | 2.12 | 627.92 | 296.36 |
| MND1 | 6.73E-10 | 1.37E-08 | -2.61 | 57.23 | 149.14 |
| CDRT1 | 9.75E-10 | 1.95E-08 | 2.90 | 149.64 | 51.53 |
| NUAK2 | 9.78E-10 | 1.96E-08 | -2.03 | 150.57 | 305.19 |
| DIO3OS | 9.84E-10 | 1.97E-08 | -2.24 | 90.32 | 202.16 |
| NALT1 | 1.99E-09 | 3.82E-08 | 2.08 | 269.06 | 129.15 |
| ASB9 | 2.35E-09 | 4.47E-08 | -2.35 | 79.36 | 186.60 |
| S1PR1 | 5.92E-09 | 1.08E-07 | 4.90 | 66.37 | 13.55 |
| PELI2 | 8.79E-09 | 1.57E-07 | 2.52 | 144.46 | 57.34 |
| SLC47A2 | 9.35E-09 | 1.67E-07 | 4.06 | 81.37 | 20.02 |
| C1orf116 | 1.37E-08 | 2.41E-07 | -2.51 | 52.58 | 131.75 |
| JAG1 | 1.55E-08 | 2.70E-07 | 2.06 | 235.18 | 114.13 |
| ETV5 | 1.69E-08 | 2.92E-07 | 3.23 | 887.53 | 274.74 |
| DUSP6 | 1.80E-08 | 3.10E-07 | 2.62 | 157.86 | 60.24 |
| LOC1001302 | 2.13E-08 | 3.60E-07 | 2.90 | 148.78 | 51.34 |
| SERTAD4 | 2.25E-08 | 3.78E-07 | -2.55 | 50.15 | 127.70 |
| LYPD3 | 2.50E-08 | 4.17E-07 | 2.77 | 126.16 | 45.48 |
| CNTD2 | 2.69E-08 | 4.49E-07 | 5.05 | 62.78 | 12.43 |
| BEND6 | 3.17E-08 | 5.25E-07 | 2.08 | 193.76 | 93.02 |
| FAM72B | 3.34E-08 | 5.52E-07 | -2.32 | 81.74 | 189.40 |
| THSD1 | 3.38E-08 | 5.56E-07 | 2.64 | 135.60 | 51.42 |
| CYGB | 3.42E-08 | 5.61E-07 | 2.67 | 136.99 | 51.32 |
| SMIM10L2A | 4.10E-08 | 6.67E-07 | 2.54 | 128.11 | 50.38 |
| EXOC3L4 | 4.17E-08 | 6.77E-07 | -3.25 | 28.76 | 93.34 |
| KLHDC7A | 4.19E-08 | 6.80E-07 | 3.18 | 89.78 | 28.27 |
| PCLAF | 4.29E-08 | 6.95E-07 | -2.24 | 1377.52 | 3091.78 |
| ZNF620 | 6.68E-08 | 1.05E-06 | -2.44 | 49.24 | 119.94 |
| DDR2 | 6.88E-08 | 1.08E-06 | 2.18 | 193.07 | 88.72 |

Supplementary Table 1: Differentially expressed genes in SiHa cells after KTI-240 treatment.

| | | | | | |
|---|---|---|---|---|---|
| COL4A3 | 7.62E-08 | 1.18E-06 | 2.17 | 197.20 | 90.84 |
| ETV4 | 7.85E-08 | 1.21E-06 | 2.61 | 250.83 | 96.23 |
| SYCE2 | 1.06E-07 | 1.59E-06 | -2.17 | 74.67 | 161.91 |
| FBXO43 | 1.40E-07 | 2.08E-06 | -2.21 | 75.30 | 166.19 |
| LOC1001299 | 1.49E-07 | 2.20E-06 | 2.27 | 233.38 | 102.68 |
| NDUFA4L2 | 1.76E-07 | 2.56E-06 | 6.47 | 42.23 | 6.53 |
| PTPRQ | 2.09E-07 | 3.00E-06 | -2.26 | 112.64 | 254.59 |
| LINC01759 | 2.54E-07 | 3.60E-06 | 2.60 | 107.13 | 41.17 |
| LOC1057476 | 3.06E-07 | 4.29E-06 | 3.33 | 79.37 | 23.81 |
| EBF1 | 3.98E-07 | 5.48E-06 | -3.58 | 17.42 | 62.31 |
| FXYD3 | 5.43E-07 | 7.30E-06 | 2.05 | 152.46 | 74.22 |
| GADD45G | 7.05E-07 | 9.21E-06 | 2.72 | 88.23 | 32.43 |
| NEAT1 | 7.49E-07 | 9.71E-06 | 2.03 | 112272.62 | 55190.41 |
| C15orf48 | 8.59E-07 | 1.10E-05 | -3.92 | 17.70 | 69.38 |
| B3GNT7 | 8.75E-07 | 1.12E-05 | -2.57 | 35.72 | 91.76 |
| TRIB3 | 9.30E-07 | 1.18E-05 | 2.94 | 3392.29 | 1153.53 |
| XKR5 | 1.04E-06 | 1.30E-05 | -2.06 | 69.08 | 142.27 |
| SSPO | 1.15E-06 | 1.43E-05 | 2.46 | 187.18 | 76.16 |
| A2MP1 | 1.61E-06 | 1.94E-05 | -2.77 | 30.45 | 84.42 |
| AKR1B10 | 1.68E-06 | 2.02E-05 | 3.02 | 110.81 | 36.73 |
| GAS6-AS1 | 1.98E-06 | 2.35E-05 | 2.79 | 77.49 | 27.78 |
| ACTBL2 | 2.01E-06 | 2.38E-05 | 5.43 | 39.82 | 7.33 |
| LDHC | 2.08E-06 | 2.45E-05 | -2.18 | 54.84 | 119.48 |
| RAB44 | 2.17E-06 | 2.54E-05 | 4.44 | 44.15 | 9.94 |
| FCGBP | 2.61E-06 | 3.00E-05 | 2.34 | 123.97 | 52.99 |
| PROC | 3.44E-06 | 3.84E-05 | 2.03 | 155.48 | 76.41 |
| DNAJC12 | 3.97E-06 | 4.36E-05 | 2.17 | 113.77 | 52.34 |
| LINC00689 | 4.04E-06 | 4.43E-05 | -2.20 | 55.49 | 121.92 |
| SFRP1 | 4.40E-06 | 4.80E-05 | -2.03 | 115.13 | 234.24 |
| FBP1 | 4.47E-06 | 4.86E-05 | -3.91 | 11.77 | 46.05 |
| HAS2 | 5.67E-06 | 6.00E-05 | -2.23 | 61.15 | 136.59 |
| CCR3 | 6.06E-06 | 6.39E-05 | -3.00 | 20.20 | 60.60 |
| SCARNA7 | 6.10E-06 | 6.42E-05 | 4.16 | 47.49 | 11.42 |
| CPB1 | 6.19E-06 | 6.51E-05 | -2.48 | 33.48 | 83.16 |
| CYP19A1 | 6.44E-06 | 6.75E-05 | 2.94 | 79.43 | 26.98 |
| C2CD4A | 6.71E-06 | 6.98E-05 | 2.91 | 66.64 | 22.87 |
| LOC729603 | 6.84E-06 | 7.10E-05 | 2.15 | 168.66 | 78.49 |
| MAFB | 7.90E-06 | 8.09E-05 | 2.64 | 108.18 | 41.02 |
| LINC00887 | 7.95E-06 | 8.13E-05 | 2.09 | 192.45 | 92.27 |
| CECR2 | 9.24E-06 | 9.32E-05 | 2.85 | 61.51 | 21.56 |
| ZNF449 | 1.23E-05 | 1.21E-04 | 2.44 | 98.61 | 40.43 |
| KANK3 | 1.26E-05 | 1.24E-04 | 2.07 | 143.30 | 69.26 |
| DDIT3 | 1.30E-05 | 1.27E-04 | 2.49 | 842.21 | 337.88 |

Supplementary Table 1:  Differentially expressed genes in SiHa cells after KTI-240 treatment.

| | | | | | |
|---|---|---|---|---|---|
| RASD1 | 1.47E-05 | 1.42E-04 | 2.02 | 135.55 | 67.13 |
| KCNK6 | 1.48E-05 | 1.43E-04 | 2.49 | 103.82 | 41.76 |
| PEG10 | 1.48E-05 | 1.43E-04 | -3.85 | 12.78 | 49.17 |
| RORA | 1.49E-05 | 1.44E-04 | 2.06 | 128.24 | 62.18 |
| KLK10 | 1.89E-05 | 1.77E-04 | -2.46 | 37.58 | 92.56 |
| POU2F3 | 1.92E-05 | 1.80E-04 | -2.33 | 35.68 | 83.28 |
| LINC00475 | 2.10E-05 | 1.94E-04 | 2.62 | 71.65 | 27.34 |
| CATSPERG | 2.17E-05 | 2.00E-04 | 2.20 | 154.27 | 70.15 |
| TMEM88 | 2.36E-05 | 2.15E-04 | 5.86 | 30.44 | 5.19 |
| VWCE | 2.58E-05 | 2.33E-04 | 3.46 | 47.69 | 13.80 |
| SCARNA9 | 3.18E-05 | 2.82E-04 | 2.71 | 105.37 | 38.84 |
| LOC1002886 | 3.62E-05 | 3.17E-04 | -2.01 | 56.23 | 112.90 |
| TMEM151A | 4.09E-05 | 3.54E-04 | 2.05 | 107.13 | 52.26 |
| CES3 | 4.13E-05 | 3.57E-04 | 2.72 | 64.91 | 23.84 |
| CYP26A1 | 4.66E-05 | 3.98E-04 | -2.70 | 21.91 | 59.07 |
| PTGS2 | 4.67E-05 | 3.98E-04 | 2.71 | 1699.92 | 627.22 |
| LSMEM1 | 4.96E-05 | 4.20E-04 | 2.67 | 56.62 | 21.23 |
| MAF | 5.30E-05 | 4.46E-04 | -2.17 | 60.22 | 130.76 |
| LUCAT1 | 5.35E-05 | 4.50E-04 | 2.50 | 79.81 | 31.92 |
| GAS6-AS2 | 5.85E-05 | 4.88E-04 | 2.71 | 57.62 | 21.30 |
| CCND1 | 7.20E-05 | 5.87E-04 | 3.09 | 61.69 | 19.99 |
| TMSB15A | 7.52E-05 | 6.10E-04 | -2.21 | 41.63 | 91.96 |
| MIR4712 | 7.77E-05 | 6.27E-04 | 3.22 | 48.52 | 15.08 |
| STC2 | 8.03E-05 | 6.45E-04 | 3.55 | 2110.22 | 594.70 |
| WNT5B | 8.43E-05 | 6.75E-04 | -2.19 | 35.42 | 77.64 |
| ELMO1 | 8.46E-05 | 6.77E-04 | -2.04 | 50.71 | 103.23 |
| CREG2 | 8.66E-05 | 6.90E-04 | -2.14 | 44.85 | 96.04 |
| KLHDC7B | 9.66E-05 | 7.61E-04 | 2.55 | 110.38 | 43.32 |
| UNC5B-AS1 | 1.13E-04 | 8.72E-04 | 4.63 | 27.54 | 5.95 |
| CAPSL | 1.16E-04 | 8.91E-04 | -3.83 | 9.60 | 36.76 |
| CEMIP | 1.70E-04 | 1.26E-03 | 3.03 | 78.37 | 25.86 |
| EDA2R | 2.02E-04 | 1.46E-03 | 3.33 | 36.00 | 10.82 |
| APOBEC3D | 2.18E-04 | 1.57E-03 | 3.22 | 48.86 | 15.16 |
| LOC1001294 | 2.23E-04 | 1.60E-03 | 2.02 | 84.57 | 41.91 |
| FAM106A | 2.26E-04 | 1.62E-03 | 2.26 | 109.50 | 48.53 |
| RRAD | 2.33E-04 | 1.66E-03 | 2.83 | 54.99 | 19.43 |
| SHC4 | 2.51E-04 | 1.77E-03 | 2.78 | 45.18 | 16.25 |
| CHRNB4 | 2.60E-04 | 1.82E-03 | -2.62 | 26.82 | 70.24 |
| FAM13A-AS1 | 2.98E-04 | 2.05E-03 | 2.73 | 54.18 | 19.86 |
| TBX2 | 3.32E-04 | 2.26E-03 | 2.42 | 57.07 | 23.55 |
| CYP26C1 | 3.47E-04 | 2.34E-03 | -3.67 | 7.96 | 29.17 |
| PDE4B | 3.69E-04 | 2.47E-03 | -2.30 | 28.62 | 65.91 |
| EP300-AS1 | 3.76E-04 | 2.51E-03 | -2.14 | 32.51 | 69.50 |

Supplementary Table 1: Differentially expressed genes in SiHa cells after KTI-240 treatment.

| | | | | | |
|---|---|---|---|---|---|
| KLF15 | 3.83E-04 | 2.55E-03 | 2.10 | 83.84 | 39.87 |
| CACNA2D3 | 3.91E-04 | 2.60E-03 | -2.22 | 30.55 | 67.73 |
| HSPA6 | 4.26E-04 | 2.79E-03 | 2.34 | 74.68 | 31.98 |
| GALNT16 | 4.40E-04 | 2.86E-03 | -2.41 | 23.94 | 57.65 |
| DIO3 | 4.43E-04 | 2.88E-03 | -3.79 | 7.34 | 27.78 |
| GCOM1 | 5.13E-04 | 3.27E-03 | 2.63 | 41.73 | 15.89 |
| HIST1H3C | 5.35E-04 | 3.40E-03 | -2.09 | 39.61 | 82.65 |
| PDE4C | 5.53E-04 | 3.49E-03 | 2.50 | 62.29 | 24.87 |
| KANK4 | 6.53E-04 | 4.04E-03 | -2.78 | 15.79 | 43.94 |
| LGI4 | 6.90E-04 | 4.25E-03 | 2.51 | 47.06 | 18.73 |
| RADIL | 7.28E-04 | 4.45E-03 | -2.09 | 32.43 | 67.81 |
| TLR6 | 7.54E-04 | 4.58E-03 | 2.28 | 55.79 | 24.49 |
| CCR1 | 8.31E-04 | 5.00E-03 | -3.64 | 8.59 | 31.23 |
| KLRA1P | 9.91E-04 | 5.79E-03 | 2.06 | 61.64 | 29.92 |
| MROH2A | 9.94E-04 | 5.80E-03 | 2.86 | 49.22 | 17.21 |
| NECAB2 | 9.95E-04 | 5.80E-03 | 2.48 | 44.61 | 18.01 |
| H2BFS | 1.00E-03 | 5.82E-03 | -2.01 | 37.26 | 74.91 |
| P2RY2 | 1.00E-03 | 5.83E-03 | -2.06 | 31.26 | 64.27 |
| KCNQ1OT1 | 1.10E-03 | 6.33E-03 | 3.84 | 1203.41 | 313.57 |
| LINC00685 | 1.14E-03 | 6.49E-03 | 2.79 | 33.76 | 12.10 |
| KRT86 | 1.21E-03 | 6.82E-03 | -2.38 | 18.38 | 43.79 |
| MAN1A1 | 1.21E-03 | 6.85E-03 | 3.26 | 29.92 | 9.17 |
| CXCL3 | 1.22E-03 | 6.89E-03 | 2.10 | 68.09 | 32.42 |
| PRKCB | 1.37E-03 | 7.63E-03 | -2.52 | 18.62 | 46.86 |
| SYNE4 | 1.37E-03 | 7.63E-03 | 2.87 | 33.93 | 11.82 |
| LOC729652 | 1.44E-03 | 7.95E-03 | 2.79 | 37.68 | 13.50 |
| MYT1L | 1.46E-03 | 8.03E-03 | -2.32 | 19.97 | 46.25 |
| GPR19 | 1.47E-03 | 8.09E-03 | -3.42 | 9.02 | 30.85 |
| MYLK3 | 1.49E-03 | 8.17E-03 | -2.23 | 22.80 | 50.94 |
| FBLN7 | 1.51E-03 | 8.29E-03 | -2.84 | 12.46 | 35.37 |
| CDHR1 | 1.67E-03 | 9.04E-03 | 3.48 | 25.35 | 7.29 |
| PYGM | 1.73E-03 | 9.30E-03 | 2.68 | 44.98 | 16.78 |
| SCARNA2 | 1.84E-03 | 9.83E-03 | 2.22 | 80.36 | 36.26 |
