## Supplementary Table 2 for "Covalent Inhibition of the Human Papillomavirus Type 16 E6 Protein Restores p53 and Suppresses HPV-Driven Tumorigenesis"

Differentially expressed genes in SiHa C51S cells after KTI-240 treatment

| Feature ID | P-value | FDR | Fold change | LSMean(KTI-240) | LSMean(DMSO) |
|---|---|---|---|---|---|
| FAM129A | 1.38E-33 | 2.18E-29 | 4.97 | 640 | 129 |
| SEC24D | 4.72E-31 | 3.73E-27 | 3.03 | 5366 | 1773 |
| GCLC | 7.47E-28 | 3.93E-24 | 2.96 | 7328 | 2474 |
| HSPA5 | 2.36E-25 | 9.31E-22 | 3.25 | 102589 | 31559 |
| SSR3 | 9.80E-25 | 3.09E-21 | 2.30 | 13306 | 5785 |
| DNAJB9 | 1.93E-23 | 5.08E-20 | 2.70 | 1949 | 722 |
| SLC12A3 | 1.92E-22 | 4.34E-19 | -5.88 | 81 | 474 |
| FAM114A1 | 2.45E-21 | 4.83E-18 | 2.80 | 1723 | 616 |
| CAMK1D | 2.33E-20 | 4.09E-17 | -2.28 | 385 | 878 |
| IGSF10 | 4.69E-20 | 7.41E-17 | -3.86 | 201 | 777 |
| AOX1 | 7.45E-20 | 1.07E-16 | -4.07 | 1249 | 5087 |
| TROAP | 2.21E-19 | 2.68E-16 | -2.04 | 1345 | 2737 |
| CALR | 4.58E-18 | 5.17E-15 | 2.60 | 70560 | 27182 |
| PSAT1 | 5.71E-18 | 6.02E-15 | 2.49 | 8880 | 3563 |
| KDELR3 | 1.43E-17 | 1.41E-14 | 2.87 | 3050 | 1063 |
| SLC7A11 | 1.56E-17 | 1.45E-14 | 4.69 | 22442 | 4787 |
| NRCAM | 2.77E-17 | 2.43E-14 | 3.59 | 521 | 145 |
| CPA4 | 4.89E-17 | 4.07E-14 | -5.25 | 48 | 250 |
| TNS3 | 5.39E-17 | 4.26E-14 | -2.06 | 3802 | 7840 |
| NMRAL2P | 4.17E-16 | 3.14E-13 | 2.78 | 578 | 208 |
| SPC24 | 5.40E-16 | 3.88E-13 | -2.66 | 468 | 1246 |
| XBP1 | 6.23E-16 | 4.28E-13 | 2.32 | 6245 | 2687 |
| CYP24A1 | 2.28E-15 | 1.44E-12 | -2.73 | 101 | 276 |
| MANF | 2.87E-15 | 1.68E-12 | 2.50 | 7081 | 2830 |
| GFPT1 | 6.10E-15 | 3.35E-12 | 2.03 | 8230 | 4049 |
| TNS4 | 6.16E-15 | 3.35E-12 | -2.16 | 3198 | 6903 |
| UGDH | 8.12E-15 | 4.28E-12 | 2.25 | 9264 | 4123 |
| KIF18B | 1.27E-14 | 6.47E-12 | -2.29 | 1626 | 3719 |
| ZNF469 | 1.79E-14 | 8.82E-12 | 3.16 | 1552 | 491 |
| STC2 | 3.48E-14 | 1.67E-11 | 4.79 | 2149 | 449 |
| HYOU1 | 3.76E-14 | 1.75E-11 | 2.30 | 17626 | 7651 |
| C11orf86 | 4.15E-14 | 1.87E-11 | -2.10 | 1579 | 3310 |
| LAMA4 | 7.74E-14 | 3.40E-11 | -2.32 | 650 | 1508 |
| MTFR2 | 9.95E-14 | 4.25E-11 | -2.19 | 214 | 470 |
| WISP2 | 2.00E-13 | 8.10E-11 | -3.92 | 43 | 170 |
| GPR87 | 2.83E-13 | 1.12E-10 | -2.84 | 262 | 743 |
| ADM2 | 3.02E-13 | 1.17E-10 | 11.41 | 396 | 35 |
| CDCA3 | 3.25E-13 | 1.22E-10 | -2.04 | 1004 | 2053 |
| TNFAIP2 | 3.78E-13 | 1.39E-10 | -2.20 | 6569 | 14452 |
| SEC11C | 1.29E-12 | 4.52E-10 | 2.01 | 1680 | 835 |
| ETNPPL | 1.94E-12 | 6.65E-10 | -2.35 | 153 | 360 |
| AURKB | 3.02E-12 | 9.93E-10 | -2.36 | 1073 | 2534 |
| SORL1 | 3.19E-12 | 1.03E-09 | -2.06 | 477 | 986 |
| GPAT3 | 3.94E-12 | 1.24E-09 | 2.75 | 963 | 351 |

Differentially expressed genes in SiHa C51S cells after KTI-240 treatment

| | | | | | |
|---|---|---|---|---|---|
| ZWINT | 4.85E-12 | 1.49E-09 | -2.04 | 2447 | 5003 |
| HMOX1 | 4.91E-12 | 1.49E-09 | 3.78 | 59984 | 15874 |
| KRT80 | 1.22E-11 | 3.31E-09 | -2.58 | 1067 | 2755 |
| TJP3 | 2.78E-11 | 7.44E-09 | -2.90 | 136 | 395 |
| E2F8 | 3.19E-11 | 8.26E-09 | -2.53 | 270 | 683 |
| SLC2A10 | 4.20E-11 | 1.04E-08 | 2.08 | 730 | 351 |
| HAS2 | 5.77E-11 | 1.38E-08 | -2.33 | 150 | 349 |
| GABRE | 8.27E-11 | 1.95E-08 | -2.17 | 604 | 1310 |
| PYCR1 | 1.07E-10 | 2.48E-08 | 3.48 | 3035 | 871 |
| OSGIN1 | 1.40E-10 | 3.20E-08 | 3.51 | 2407 | 685 |
| RRM2 | 2.07E-10 | 4.60E-08 | -2.22 | 10364 | 22977 |
| PLPP5 | 2.12E-10 | 4.66E-08 | 2.27 | 2165 | 955 |
| A2MP1 | 2.66E-10 | 5.68E-08 | -3.76 | 33 | 125 |
| HJURP | 3.07E-10 | 6.38E-08 | -2.02 | 1010 | 2040 |
| HSP90B1 | 3.23E-10 | 6.50E-08 | 2.82 | 84577 | 29981 |
| GABRQ | 3.25E-10 | 6.50E-08 | -2.17 | 127 | 276 |
| MYBL2 | 3.46E-10 | 6.84E-08 | -2.02 | 3874 | 7829 |
| SRXN1 | 6.88E-10 | 1.29E-07 | 2.94 | 16019 | 5447 |
| CIT | 8.08E-10 | 1.48E-07 | -2.24 | 2329 | 5226 |
| ARHGEF39 | 9.60E-10 | 1.69E-07 | -2.38 | 222 | 527 |
| PIF1 | 1.11E-09 | 1.92E-07 | -2.29 | 412 | 944 |
| DHRS3 | 1.30E-09 | 2.24E-07 | -3.38 | 215 | 728 |
| KRTAP7-1 | 1.35E-09 | 2.26E-07 | -2.21 | 176 | 388 |
| PCOLCE2 | 1.47E-09 | 2.44E-07 | 2.28 | 249 | 109 |
| GTSE1 | 1.54E-09 | 2.50E-07 | -2.13 | 1679 | 3572 |
| TXNIP | 1.76E-09 | 2.83E-07 | -2.69 | 2509 | 6748 |
| STEAP4 | 2.34E-09 | 3.69E-07 | -2.57 | 20451 | 52551 |
| WIPI1 | 4.13E-09 | 6.28E-07 | 2.26 | 1644 | 728 |
| MAF | 4.13E-09 | 6.28E-07 | -2.28 | 185 | 422 |
| NAV2 | 4.54E-09 | 6.77E-07 | -2.13 | 859 | 1827 |
| CHAC1 | 4.95E-09 | 7.30E-07 | 4.43 | 525 | 118 |
| RAD54L | 6.05E-09 | 8.77E-07 | -2.35 | 563 | 1323 |
| PSRC1 | 9.73E-09 | 1.36E-06 | -2.09 | 525 | 1098 |
| PPL | 1.48E-08 | 2.01E-06 | -2.09 | 4913 | 10290 |
| SLC1A5 | 1.51E-08 | 2.04E-06 | 2.62 | 19635 | 7488 |
| NCAPH | 3.06E-08 | 3.96E-06 | -2.27 | 948 | 2152 |
| TRIB3 | 4.20E-08 | 5.31E-06 | 4.09 | 2796 | 684 |
| PHGDH | 4.39E-08 | 5.50E-06 | 2.53 | 7028 | 2781 |
| MXD3 | 4.45E-08 | 5.50E-06 | -2.32 | 362 | 841 |
| LINC01260 | 4.46E-08 | 5.50E-06 | -7.81 | 5 | 41 |
| CXCL1 | 5.74E-08 | 6.87E-06 | -2.70 | 141 | 379 |
| FZD10 | 6.73E-08 | 7.87E-06 | -2.38 | 50 | 118 |
| PRKCB | 7.17E-08 | 8.33E-06 | -3.26 | 23 | 76 |
| TIMP3 | 8.76E-08 | 9.96E-06 | -2.06 | 2364 | 4865 |
| EBF1 | 9.75E-08 | 1.09E-05 | -6.19 | 6 | 35 |

Differentially expressed genes in SiHa C51S cells after KTI-240 treatment

| | | | | | |
|---|---|---|---|---|---|
| MYT1L | 1.50E-07 | 1.59E-05 | -2.51 | 83 | 209 |
| DDX12P | 1.51E-07 | 1.59E-05 | -2.08 | 671 | 1400 |
| VSIR | 1.62E-07 | 1.68E-05 | -2.46 | 538 | 1326 |
| ZBED6CL | 1.62E-07 | 1.68E-05 | -2.09 | 872 | 1826 |
| KIF20A | 1.80E-07 | 1.82E-05 | -2.23 | 4297 | 9603 |
| FBN1 | 2.02E-07 | 1.98E-05 | -2.88 | 25 | 73 |
| TNFSF10 | 2.18E-07 | 2.09E-05 | -4.01 | 36 | 146 |
| MYLK2 | 2.52E-07 | 2.36E-05 | -2.58 | 46 | 118 |
| LINC00689 | 2.72E-07 | 2.48E-05 | -2.88 | 32 | 91 |
| EPGN | 3.29E-07 | 2.96E-05 | 2.89 | 3323 | 1149 |
| TNFSF14 | 3.55E-07 | 3.15E-05 | -4.97 | 12 | 59 |
| CRISPLD2 | 3.72E-07 | 3.28E-05 | 2.06 | 1876 | 911 |
| DKK1 | 4.37E-07 | 3.76E-05 | -2.10 | 3930 | 8250 |
| PPIB | 4.37E-07 | 3.76E-05 | 2.21 | 43775 | 19849 |
| BDKRB1 | 4.51E-07 | 3.85E-05 | -2.64 | 50 | 132 |
| PDGFRL | 5.22E-07 | 4.36E-05 | 3.09 | 255 | 83 |
| AQP3 | 6.90E-07 | 5.61E-05 | -2.05 | 580 | 1187 |
| BHLHE41 | 7.43E-07 | 5.96E-05 | 2.40 | 113 | 47 |
| SLC6A9 | 8.56E-07 | 6.76E-05 | 2.38 | 2554 | 1074 |
| ARL14EPL | 8.61E-07 | 6.77E-05 | -2.51 | 59 | 148 |
| VEPH1 | 8.90E-07 | 6.93E-05 | -2.21 | 48 | 105 |
| ANXA8 | 9.10E-07 | 7.01E-05 | -2.14 | 134 | 288 |
| JPH2 | 9.60E-07 | 7.25E-05 | -2.03 | 121 | 246 |
| ARHGAP33 | 1.05E-06 | 7.81E-05 | -2.05 | 379 | 778 |
| PTPRQ | 1.23E-06 | 8.98E-05 | -2.82 | 51 | 143 |
| APBB1IP | 1.28E-06 | 9.31E-05 | -2.23 | 255 | 568 |
| FOSL1 | 1.72E-06 | 1.19E-04 | 2.30 | 1139 | 496 |
| ATF3 | 1.84E-06 | 1.26E-04 | 2.37 | 622 | 263 |
| RIPOR2 | 1.89E-06 | 1.28E-04 | -2.57 | 35 | 90 |
| NREP | 1.90E-06 | 1.29E-04 | -2.19 | 274 | 598 |
| MUC16 | 2.66E-06 | 1.69E-04 | -3.16 | 399 | 1263 |
| LOC1002886 | 2.82E-06 | 1.77E-04 | -2.05 | 65 | 134 |
| PCLAF | 3.14E-06 | 1.93E-04 | -2.07 | 2404 | 4975 |
| NRP2 | 3.19E-06 | 1.96E-04 | -2.91 | 47 | 137 |
| MTUS1 | 3.57E-06 | 2.16E-04 | -2.35 | 274 | 643 |
| EBF2 | 3.84E-06 | 2.32E-04 | 2.43 | 103 | 42 |
| CNTD2 | 4.14E-06 | 2.47E-04 | 6.49 | 42 | 6 |
| NRGN | 4.42E-06 | 2.60E-04 | -2.63 | 35 | 93 |
| C1QTNF2 | 4.88E-06 | 2.83E-04 | -4.62 | 8 | 36 |
| UCHL1 | 5.60E-06 | 3.20E-04 | 2.06 | 625 | 303 |
| TK1 | 5.95E-06 | 3.37E-04 | -2.13 | 1731 | 3685 |
| PCK2 | 6.81E-06 | 3.80E-04 | 2.90 | 3923 | 1353 |
| NUCB2 | 8.11E-06 | 4.37E-04 | 2.35 | 5388 | 2297 |
| ASNS | 8.34E-06 | 4.45E-04 | 3.05 | 152 | 50 |
| STARD13 | 9.04E-06 | 4.81E-04 | -2.01 | 175 | 352 |

Differentially expressed genes in SiHa C51S cells after KTI-240 treatment

| | | | | | |
|---|---|---|---|---|---|
| GJA5 | 9.34E-06 | 4.95E-04 | -2.51 | 47 | 119 |
| PANX2 | 1.01E-05 | 5.33E-04 | 2.31 | 1449 | 629 |
| P4HA3 | 1.40E-05 | 7.06E-04 | 3.37 | 41 | 12 |
| ASB9 | 1.74E-05 | 8.68E-04 | -2.09 | 49 | 103 |
| GRB7 | 2.16E-05 | 1.06E-03 | -2.28 | 171 | 390 |
| VAV3 | 2.36E-05 | 1.14E-03 | -2.65 | 79 | 210 |
| FAM167A | 2.51E-05 | 1.20E-03 | 2.24 | 105 | 47 |
| LRP2 | 2.71E-05 | 1.28E-03 | -4.83 | 13 | 64 |
| OVOS | 3.13E-05 | 1.44E-03 | -2.06 | 75 | 156 |
| EHF | 3.22E-05 | 1.47E-03 | -4.30 | 15 | 66 |
| HMGB2 | 3.27E-05 | 1.49E-03 | -2.01 | 4120 | 8290 |
| UCA1 | 3.52E-05 | 1.57E-03 | -3.71 | 29 | 108 |
| TMC5 | 3.66E-05 | 1.62E-03 | -4.09 | 9 | 37 |
| SEMA3C | 4.21E-05 | 1.84E-03 | -2.18 | 3212 | 7007 |
| NCAPG | 4.24E-05 | 1.84E-03 | -2.13 | 2904 | 6200 |
| CDHR1 | 4.39E-05 | 1.90E-03 | 3.97 | 34 | 9 |
| LOC1001304 | 4.92E-05 | 2.09E-03 | -2.42 | 169 | 409 |
| GCLM | 5.34E-05 | 2.25E-03 | 2.11 | 15471 | 7323 |
| DDIT3 | 5.86E-05 | 2.43E-03 | 2.32 | 811 | 350 |
| CYP19A1 | 5.87E-05 | 2.43E-03 | 2.16 | 445 | 206 |
| SPC25 | 6.60E-05 | 2.66E-03 | -2.39 | 48 | 114 |
| BMPER | 6.84E-05 | 2.73E-03 | -2.40 | 28 | 66 |
| SERTAD4 | 7.86E-05 | 3.09E-03 | -2.16 | 58 | 125 |
| PDE5A | 1.08E-04 | 4.05E-03 | -2.35 | 656 | 1545 |
| ANGPTL4 | 1.10E-04 | 4.09E-03 | 2.36 | 122 | 52 |
| NOTUM | 1.13E-04 | 4.19E-03 | -2.20 | 389 | 857 |
| PLCL1 | 1.14E-04 | 4.22E-03 | -2.63 | 48 | 127 |
| FKBP11 | 1.27E-04 | 4.65E-03 | 2.02 | 1444 | 715 |
| MKI67 | 1.47E-04 | 5.24E-03 | -2.28 | 11711 | 26685 |
| MVD | 1.55E-04 | 5.52E-03 | 2.08 | 7747 | 3719 |
| TSHZ2 | 1.84E-04 | 6.38E-03 | -2.01 | 308 | 620 |
| EIF4EBP1 | 1.88E-04 | 6.42E-03 | 2.38 | 1670 | 702 |
| FAP | 1.99E-04 | 6.76E-03 | -2.16 | 151 | 326 |
| ECM2 | 2.09E-04 | 7.05E-03 | 3.83 | 27 | 7 |
| ANLN | 2.55E-04 | 8.34E-03 | -2.18 | 6675 | 14540 |
| MMRN2 | 2.83E-04 | 9.08E-03 | -2.24 | 28 | 64 |
| MFSD2A | 3.09E-04 | 9.68E-03 | 2.04 | 495 | 243 |
| LCP1 | 3.12E-04 | 9.71E-03 | 3.30 | 31 | 9 |
| CMAHP | 3.16E-04 | 9.83E-03 | -3.08 | 11 | 34 |
