## Supplementary Table 3 for "Covalent Inhibition of the Human Papillomavirus Type 16 E6 Protein Restores p53 and Suppresses HPV-Driven Tumorigenesis"

Differentially expressed genes in RPE-1 cells after KTI-240 treatment

| Feature ID | P-value | FDR | Fold change | LSMean(KTI-240) | LSMean (DMSO) |
|---|---|---|---|---|---|
| DHRS3 | 4.05E-51 | 6.03E-47 | -3.28 | 3301.6 | 10825.6 |
| SLC7A11 | 6.60E-35 | 4.91E-31 | 4.10 | 7227.3 | 1763.6 |
| VSIR | 1.48E-25 | 7.34E-22 | -2.49 | 342.6 | 853.3 |
| NANOS1 | 1.03E-23 | 3.83E-20 | -2.94 | 155.2 | 456.0 |
| GPRC5A | 1.59E-23 | 4.73E-20 | -2.03 | 10437.3 | 21137.1 |
| MIR210HG | 1.69E-15 | 2.52E-12 | -3.31 | 99.2 | 328.4 |
| NEAT1 | 1.26E-14 | 1.56E-11 | 2.35 | 45057.6 | 19139.6 |
| GPRC5B | 1.18E-13 | 1.17E-10 | -2.34 | 987.7 | 2310.5 |
| FAM20C | 1.51E-11 | 1.11E-08 | -2.43 | 3149.4 | 7660.5 |
| NFIA-AS2 | 6.88E-10 | 3.66E-07 | -2.79 | 73.1 | 203.7 |
| OSGIN1 | 1.32E-09 | 6.78E-07 | 2.16 | 545.6 | 252.6 |
| NMRAL2P | 4.69E-09 | 2.11E-06 | 2.95 | 173.8 | 58.9 |
| ANKRD37 | 9.15E-09 | 3.68E-06 | -2.09 | 108.1 | 225.9 |
| GCLM | 1.51E-08 | 5.46E-06 | 2.18 | 1790.7 | 820.4 |
| BST2 | 9.44E-08 | 2.30E-05 | 2.21 | 207.2 | 93.6 |
| IFIT2 | 2.19E-07 | 4.79E-05 | 2.30 | 2435.7 | 1059.2 |
| PPFIA4 | 1.65E-06 | 2.52E-04 | -3.31 | 46.0 | 152.5 |
| ARL14EPL | 8.20E-06 | 8.92E-04 | -3.50 | 15.8 | 55.3 |
| IFI44L | 8.26E-06 | 8.92E-04 | 2.12 | 1531.6 | 722.2 |
| MX2 | 1.24E-05 | 1.25E-03 | 6.30 | 106.2 | 16.8 |
| LINC00857 | 1.41E-05 | 1.39E-03 | -2.15 | 75.8 | 162.5 |
| CYP26B1 | 1.52E-05 | 1.49E-03 | -2.90 | 32.2 | 93.3 |
| MYB | 7.73E-05 | 5.48E-03 | -2.53 | 36.4 | 92.0 |
| RARA-AS1 | 8.73E-05 | 6.10E-03 | -2.03 | 67.9 | 137.9 |
| PGM5P2 | 1.48E-04 | 8.76E-03 | 2.11 | 434.5 | 206.0 |
